# Mitochondrial carrier SLC25A34 links clock, diet, and temperature control of interorganellar lipid cycling

**DOI:** 10.64898/2026.05.30.724257

**Authors:** Iuliia Karavaeva, Astrid Linde Basse, Samuel A. J. Trammell, Mohammed Faiz Hussain, Lasse Kruse Markussen, Jesper F. Havelund, Marie Sophie Isidor, Hannah J. Richter, Sabina Chubanava, Yann Deleye, Zafir Kaiser, Yachen Shen, Ditte Neess, Frederike Sass, Fabian Finger, Lidia Argemi Muntadas, David Tandio, Tao Ma, Elahu Gosney Sustarsic, Hannes Embring, Cecilie Kynding Kristensen, Rebecca L. McIntyre, Genesee J. Martinez, Anna Sofie Husted, Matthew J. Emmett, Zachary A. Kipp, Mikkel Frost, Mark P. Jedrychowski, Michel van Weeghel, Homa Majd, Ekaterina Zhuravleva, Robert W. McGarrah, Kaja Plucinska, Mohit K. Midha, Andreas Prokesch, Paul Cohen, James G. Granneman, Patrick Seale, Riekelt H. Houtkooper, Jacob B. Hansen, Steven P. Gygi, Thue W. Schwartz, Matthew Paul Gillum, Terry D. Hinds, Raymond Edward Soccio, Phillip J. White, Edmund R. S. Kunji, Thomas Moritz, Jonas T. Treebak, Susanne Mandrup, Brice Emanuelli, Lawrence Kazak, Nils J. Færgeman, Mitchell A. Lazar, Zachary Gerhart-Hines

**Affiliations:** Novo Nordisk Foundation Center for Basic Metabolic Research, University of Copenhagen, Copenhagen, DK; Center for Adipocyte Signaling (ADIPOSIGN), University of Southern Denmark, Odense, DK; Department of Biomedical Sciences, University of Copenhagen, Copenhagen, DK; Department of Biochemistry, Goodman Cancer Research Centre, McGill University Montreal, Quebec, CA; Functional Genomics & Metabolism Research Unit, Department of Biochemistry and Molecular Biology, University of Southern Denmark, Odense, DK; Department of Biochemistry and Molecular Biology, University of Southern Denmark, Odense, DK; Institute for Diabetes, Obesity, and Metabolism, University of Pennsylvania Perelman School of Medicine, Philadelphia, PA, USA; Division of Endocrinology, Diabetes, and Metabolism, Department of Medicine, University of Pennsylvania Perelman School of Medicine, Philadelphia, PA, USA; Gladstone Institute of Data Science and Biotechnology, Gladstone Institutes, San Francisco, CA, USA; Institute for Diabetes and Obesity, Helmholtz Diabetes Center, Munich, Germany; Sarah W. Stedman Nutrition and Metabolism Center and Duke Molecular Physiology Institute, Duke University Medical Center, Divisions of Endocrinology and Cardiology, Department of Medicine, and Department of Pharmacology and Cancer Biology, Durham, North Carolina, USA; Department of Pathology and Laboratory Medicine, University of Pennsylvania Perelman School of Medicine, Philadelphia, PA, USA; Drug & Disease Discovery D3 Research Center, Department of Pharmacology and Nutritional Sciences, University of Kentucky College of Medicine, Lexington, KY, USA; Broad Institute of MIT and Harvard, Cambridge, MA, USA; Massachusetts General Hospital Cancer Center and Department of Medicine, Harvard Medical School, Boston, USA; Department of Medical Oncology, Dana-Farber Cancer Institute, Harvard Medical School, Boston, MA, USA; Department of Cell Biology, Harvard Medical School, Boston, MA, USA; Laboratory Genetic Metabolic Diseases, Amsterdam Gastroenterology and Metabolism; Amsterdam UMC, University of Amsterdam, Amsterdam, NL; Medical Research Council Mitochondrial Biology Unit, University of Cambridge, Cambridge Biomedical Campus, Cambridge, UK; LEO Foundation Skin Immunology Research Center, Department of Immunology and Microbiology, Faculty of Health and Medical Sciences, University of Copenhagen, DK; Laboratory of Molecular Metabolism, The Rockefeller University, New York, New York, USA; Division of Cell Biology, Histology and Embryology, Gottfried Schatz Research Center for Cell Signaling, Metabolism and Aging, Medical University of Graz, Graz, AT; Center for Molecular Medicine and Genetics, Wayne State University School of Medicine, Detroit, MI, USA; Department of Biology, University of Copenhagen, Copenhagen, DK

## Abstract

Adipocyte lipid metabolism is coordinated by circadian rhythms, diet, and environmental temperature. Yet how these diverse signals are molecularly integrated remains unknown. Here we show that clock, diet, and temperature cues converge on the orphan mitochondrial transporter, SLC25A34, to orchestrate thermogenic cycling of lipid synthesis and oxidation. During sleep, the clock suppresses *Slc25a34* transcription through REV-ERBα. Waking, lipid-rich diets, or cold exposure abolish this repression, allowing lipolytic signals to stimulate *Slc25a34* expression via PPARα. SLC25A34 then imports oxaloacetate into mitochondria to accelerate the export of substrates used for acetyl-CoA production in the cytosol. This feeds into cytosolic lipid synthesis and transcriptional induction of mitochondrial biogenesis, which collectively promote mitochondrial lipid oxidation. Thus, SLC25A34 confers circadian, dietary, and environmental control of thermogenic metabolism through interorganellar lipid cycling.

## MAIN TEXT

The circadian clock is responsible for synchronizing physiological processes with environmental cues. This intricate system is present in nearly every cell of the body and integrates signals from sunlight, diet, physical activity, and environmental temperature to optimize metabolic function across tissues (*1–3*). One of the challenges in circadian metabolic control is the need to maintain consistent daily oscillations in anticipation of expected energetic needs while also facilitating the ability to acutely respond and adapt to unexpected perturbations.

Adipose thermogenesis is an ideal paradigm in which to study this coordinated anticipatory and adaptive nature of the clock. The energy-expending activity of thermogenic adipocytes is regulated by exposure to cold temperature (*4–6*), diet composition and quantity consumed (*7*, *8*), and the daily oscillations of the circadian clock (*9–11*) in mice and humans. In rodents, these energy-dissipating cells are critical for body temperature defense (*12*), whereas in humans, brown adipose tissue (BAT) appears to improve lipid and glucose homeostasis and is associated with cardiometabolic protection (*13*). In mice and humans, the circadian rhythm of BAT activity begins to increase just before waking (*9*, *14*, *15*) and may be involved in triggering arousal from sleep. It reaches its peak late in the waking period and then begins to drop until its lowest point during sleep. Given the energetic cost of BAT activity and the caloric scarcity our ancestors faced, clock repression of thermogenesis during sleep may have conferred an evolutionary advantage to reduce unnecessary energy expenditure.

Yet, regardless of time of day, acute exposure to cold temperature can override this circadian control and rapidly stimulate adipose thermogenesis through the sympathetic nervous system (*4*). While the specific metabolic pathways involved in the interaction between clock and temperature cues are unresolved, lipid utilization likely plays a role as numerous elements of adipocyte lipid metabolism are regulated by both programs (*4*, *16*, *17*). Notably, a unique hallmark of BAT lipid metabolism is the ability to simultaneously increase lipid synthesis in the cytosol and fatty acid oxidation in the mitochondria (*18–20*). This paradigm of synthesis and breakdown fuels an energy-dissipating, interorganellar lipid cycle that physiologically promotes thermogenesis and has more recently gained interest as a promising therapeutic strategy to counteract the caloric excess of obesity and metabolic diseases (*21*, *22*). Both environmental cold (*18–20*) and the circadian clock (*23*) have been individually linked to influencing this dual activation of anabolic and catabolic lipid pathways. Yet, how continuous circadian rhythmicity and acute cold responsiveness are molecularly integrated to coordinate thermogenic metabolism remains unknown.

### Clock and temperature control of the orphan mitochondrial transporter, *Slc25a34*

To understand how circadian and cold-regulated signaling pathways orchestrate thermogenic metabolism in adipocyte, we employed a comparative, multi-omics approach to identify candidate factors in BAT using the following criteria (**Fig. 1A, Table S1**): (I) proteins that were induced by cold (4°C cold exposure for 3 weeks vs thermoneutrality, Fold Change > 2) (*24*), (II) proteins increased/de-repressed by *Rev-erbα* genetic deletion (*Rev-erbα* whole body knockout vs control, Fold Change > 2) (**Table S2**), (III) genes with a *Rev-erbα* binding site associated 10 kb of nearest gene with higher occupancy at thermoneutrality vs cold (*9*), and (IV) genes that were regulated both by cold and REV-ERBα/β at the transcriptional level (FC >2 in wild-type mice upon cold exposure and in Rev-erbα/β double knockout vs. wild-type mice, FDR < 0.05 in all cases) (**Table S3**). Only two factors satisfied all criteria: the well-established, canonical thermogenic effector, UCP1 (a.k.a. SLC25A7), and the uncharacterized, orphan mitochondrial metabolite transporter, SLC25A34. Notably, both proteins are members of the SLC25A family of mitochondrial solute carriers (*25*, *26*) highlighting the critical importance of metabolite and ion exchange across the inner mitochondrial membrane (IMM) for adipocyte function.

**Figure 1.**
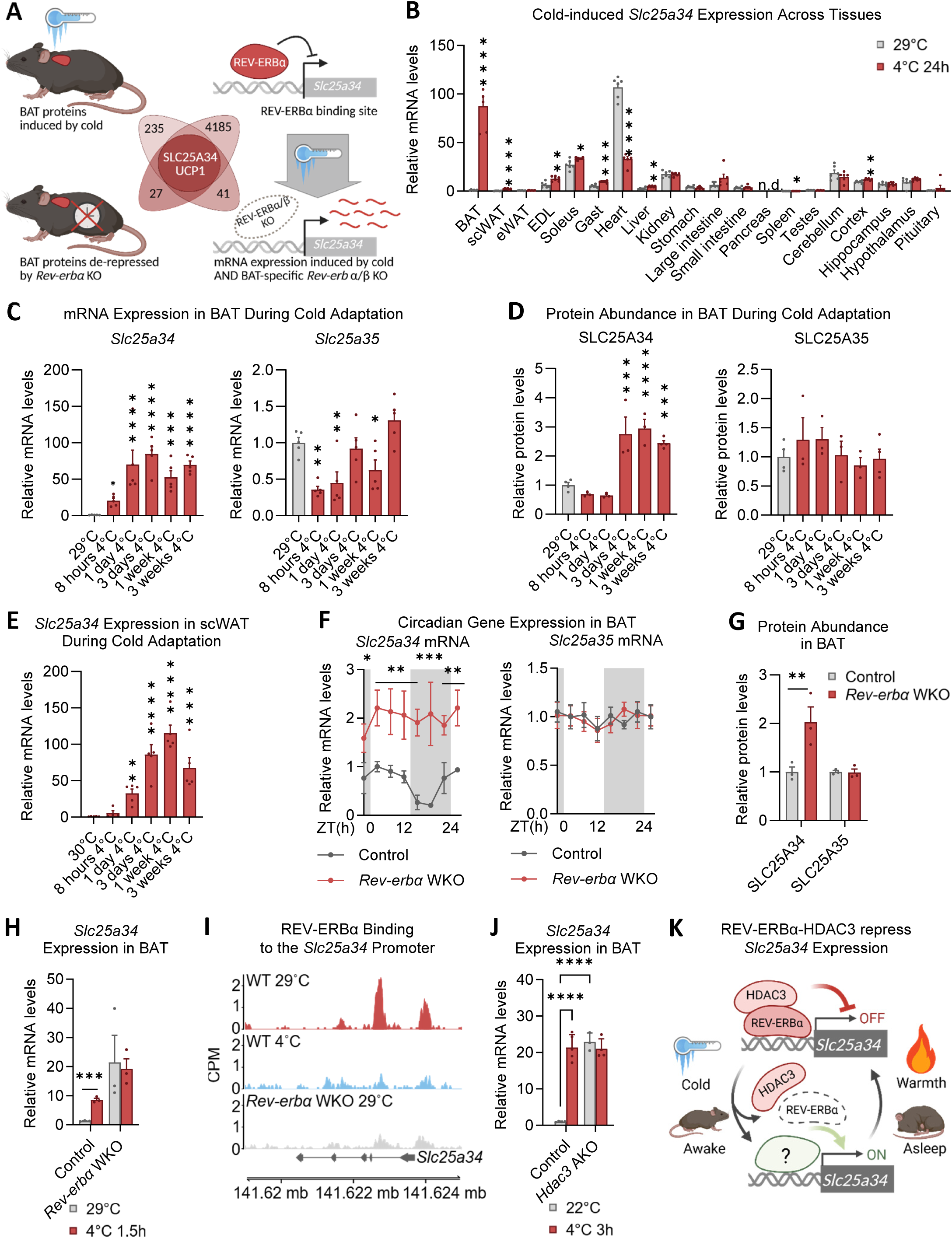
Clock and temperature control of the orphan mitochondrial transporter, *Slc25a34* via the circadian repressor, REV-ERBα. **A,** schematic of the multi-omic approach to discover *Slc25a34*; comparing the proteome of BAT from (1) cold-exposed mice (top left, 3 weeks cold exposure vs thermoneutrality, FC>2) and (2) *Rev-erbα* WKO mice (bottom left, *Rev-erbα* WKO vs Control, FC>2), (3) cistrome of BAT from Control and Rev-erbα WKO mice (top right, REV-ERBα binding sites located within 10 kb of the nearest gene that show higher occupancy in cold conditions than at thermoneutrality), and (4) BAT transcriptome from cold-exposed *Rev-erbα/β* dBKO mice (bottom right, genes induced by cold in control mice (FC > 2, FDR < 0.05) and genes induced by double Rev-erbα/β knockout in BAT at thermoneutrality (FC > 2, FDR < 0.05). **B**, *Slc25a34* expression across tissues, fold change relative to the expression level in the tissue at 29°C. **C**, *Slc25a34* and *Slc25a35* mRNA and, **D**, protein levels in BAT during cold adaptation, significance indicates effects of cold exposure compared to 29°C. **E**, *Slc25a34* mRNA levels in scWAT during cold adaptation, significance indicates effects of cold exposure compared to 29°C. **F**, Circadian expression of *Slc25a34* and *Slc25a35* mRNA levels in control and *Rev-erbα* WKO mice acclimated to 29°C. **G**, SLC25A34 and SLC25A35 protein in BAT of control and *Rev-erbα* WKO mice at 22°C at ZT10. **H**, *Slc25a34* mRNA levels in BAT of control and *Rev-erbα* WKO mice at 29°C and after 1.5h of 4°C exposure (ZT4 – ZT5.5). **I**, REV-ERBα occupancy in the proximal *Slc25a34* promoter in BAT at 29°C and after 6 h of 4°C exposure of control and *Rev-erbα* WKO mice (ZT10). **J**, *Slc25a34* mRNA levels in BAT from control and *Hdac3* AKO mice at 22°C and after 3h at 4°C. **K**, Schematic depicting the role of REV-ERBα in coordinating circadian and cold-mediated regulation of *Slc25a34* expression. For all panels, data are represented as mean ±SEM, p < 0.05 = *, p < 0.01 = **, p < 0.001 = ***, p < 0.0001 = ****, unpaired two-tailed multiple t-tests (B, H), One-way ANOVA with the Benjamini, Krieger, and Yekutieli two-stage linear step-up procedure for multiple comparisons correction (C-E), Two-way ANOVA with the Benjamini, Krieger, and Yekutieli two-stage linear step-up procedure for multiple comparisons correction (F, G, J).

At thermoneutrality, *Slc25a34* expression was the highest in the heart, followed by slow-twitch skeletal muscle, kidney, and multiple brain regions (**Fig. 1B**). BAT was among the organs with the lowest *Slc25a34* mRNA. However, upon 24 hours of cold exposure, *Slc25a34* was robustly induced in BAT (90-fold above thermoneutrality), which supplanted heart as the most *Slc25a34*-enriched tissue (**Fig. 1B**). *Slc25a34* expression was induced in BAT after only 8 hours of cold exposure and remained elevated throughout three weeks of cold adaptation (**Fig. 1C**). This was reflected at the protein level where SLC25A34 was significantly increased from 3 days of cold exposure and for the duration of cold acclimation (**Fig. 1D**). No cold induction was observed for SLC25A35, the closest homolog of SLC25A34 and which has been proposed to have a similar transport function based on sequence conservation in key substrate-binding residues (*27*, *28*). In fact, *Slc25a35* was acutely suppressed by cold exposure (**Fig. S1A, 1C**) with protein levels remaining unchanged throughout cold acclimation (**Fig. 1D**). In subcutaneous white adipose tissue (scWAT), which undergoes recruitment of energy-expending beige adipocytes during cold adaptation, both transporters were cold-induced but *Slc25a34* to a nearly 40-fold greater extent (**Fig. 1E, Fig. S1B**). Thus, the orphan mitochondrial transporter, SLC25A34, is robustly and sustainably increased in thermogenic adipose tissues following exposure to environmental cold temperature.

### Circadian and cold control of *Slc25a34* by REV-ERBα and HDAC3-mediated transcriptional repression

In addition to regulation by environmental temperature, BAT *Slc25a34* expression also robustly oscillates over 24 hours (**Fig. 1F**). Whole-body knockout of the circadian nuclear receptor *Rev-erbα* (hereby referred to as *Rev-erbα* WKO) not only abolished this rhythmicity but resulted in constitutively increased *Slc25a34* expression compared to control littermates (**Fig. 1F**), demonstrating that REV-ERBα-mediated repression was responsible for clock control of *Slc25a34*. Consistent with this, SLC25A34 protein levels were elevated in BAT from *Rev-erbα* WKO mice (**Fig. 1G**). Conversely, *Slc25a35* expression was not circadian, nor were its protein levels changed upon *Rev-erbα* deletion (**Fig. 1F, 1G**).

Given that the circadian clock, through REV-ERBα, plays a critical role in temperature-dependent thermogenic control (*9*), we investigated the interaction between REV-ERBα-suppression and cold induction of *Slc25a34*. We measured *Slc25a34* and *Slc25a35* expression in BAT from thermoneutral-acclimated and cold-exposed *Rev-erbα* WKOs and control littermates. Cold exposure increased *Slc25a34* expression in the control mice in agreement with our previous experiments (**Fig. 1B and 1C**). However, in *Rev-erbα* WKO BAT, the levels of *Slc25a34* were already maximally elevated at thermoneutrality and could not be further increased upon cold exposure (**Fig. 1H**), suggesting that de-repressing REV-ERBα control is a critical step in cold induction of the transporter. Indeed, we found that REV-ERBα binds to two sites in the proximal promoter and first intron of *Slc25a34* under thermoneutral conditions and this binding is abolished by cold temperature (**Fig. 1I**). In contrast, *Slc25a35* was downregulated by cold to the same extent in both *Rev-erbα* WKOs and control littermates **(Fig. S1C)** despite the presence of REV-ERBα binding site in its proximal promoter (**Fig. S1D**).

To determine if the observed regulation of *Slc25a34* expression in BAT was cell-autonomous and not due to central effects of whole-body *Rev-erbα* disruption, we measured expression of the transporters in brown adipocyte-specific *Rev-erbα and Rev-erbβ* double knockout mice (*Rev-erbα/β* dBKO) housed at thermoneutrality or cold-challenged (**Table S3**). *Rev-erbβ* was knocked out because it has been shown to compensate for loss of *Rev-erbα* in other tissues (*29*). *Slc25a34* expression was de-repressed in *Rev-erbα/β* dBKO mice compared to littermate controls both at thermoneutrality and in cold (**Fig. S1F**), indicating that REV-ERBs regulate *Slc25a34* expression cell-autonomously. Conversely, *Slc25a35* expression was not affected by *Rev-erbα and Rev-erbβ* double knockout at thermoneutrality and was significantly decreased by cold exposure in both genotypes (**Fig. S1F**).

REV-ERBα represses transcription through the recruitment of HDAC3 and subsequent histone deacetylation (*30*). To address whether HDAC3 contributes to cold regulation of *Slc25a34*, we measured transporter expression in the BAT of adipose-specific *Hdac3* knockout mice (*Hdac3* AKO) and littermate controls at room temperature and following an acute cold challenge. Similar to the *Rev-erbα* WKO mice, BAT *Slc25a34* expression in *Hdac3* AKO mice was elevated at room temperature in comparison to control mice and could not be further induced by cold exposure (**Fig. 1J**), indicating that HDAC3 acts as the REV-ERBα-directed corepressor of *Slc25a34*. Conversely, BAT *Slc25a35* expression was decreased in the *Hdac3* AKO in comparison to control animals at room temperature and was even further suppressed by cold in the *Hdac3* AKO mice (**Fig. S1D**). Altogether, these data show that REV-ERBα and HDAC3 coordinate the circadian and temperature-dependent regulation of *Slc25a34* in BAT by suppressing its expression at thermoneutrality and during sleep. Waking or acute cold exposure abolishes this repression and increases the abundance of the transporter (**Fig. 1K**).

### Cold and diet-induced lipid metabolism stimulate *Slc25a34* expression via PPARα

While our findings establish REV-ERBα-dependent repression to be a key event in the regulation of *Slc25a34*, the complementary transcription factor or factors mediating activation of *Slc25a34* expression remained unknown. Given the numerous transcriptional regulators of thermogenic metabolism in adipocytes (*31*), we sought to identify potential activators by performing pathway enrichment analysis on the top 50 BAT proteins that were expressed in a similar temporal pattern to SLC25A34 throughout cold adaptation (*24*) (**Fig. 2A, Table S4**). Strikingly, these proteins were nearly all linked to biological processes related to lipid metabolism and biosynthesis (**Fig. 2B**). This observation suggested that PPARs, the canonical regulators of adipocyte lipid metabolism (*32*), could be transcriptional activators of *Slc25a34*. Indeed, both PPARα and PPARγ bind to a regulatory region near *Slc25a34* that is enriched for active histone marks (*33*). Agonists for PPARα (GW7647) or PPARγ (rosiglitazone) cell autonomously induced *Slc25a34* expression in brown adipocytes alongside their canonical targets *Ucp1* and *Elovl3* (**Fig. 2C-2F, S2A, S2B**). We further found that siRNAs targeting either factor significantly decreased the mRNA levels of the transporter (**Fig. 2G, 2H**). These studies suggest that both PPARα and PPARγ regulate *Slc25a34* expression *in vitro*. However, determining specific PPAR dependency for a particular gene is confounded by the ability of PPARγ to both control PPARα expression (*34*) and compensate for *Pparα* loss (*35*).

**Figure 2.**
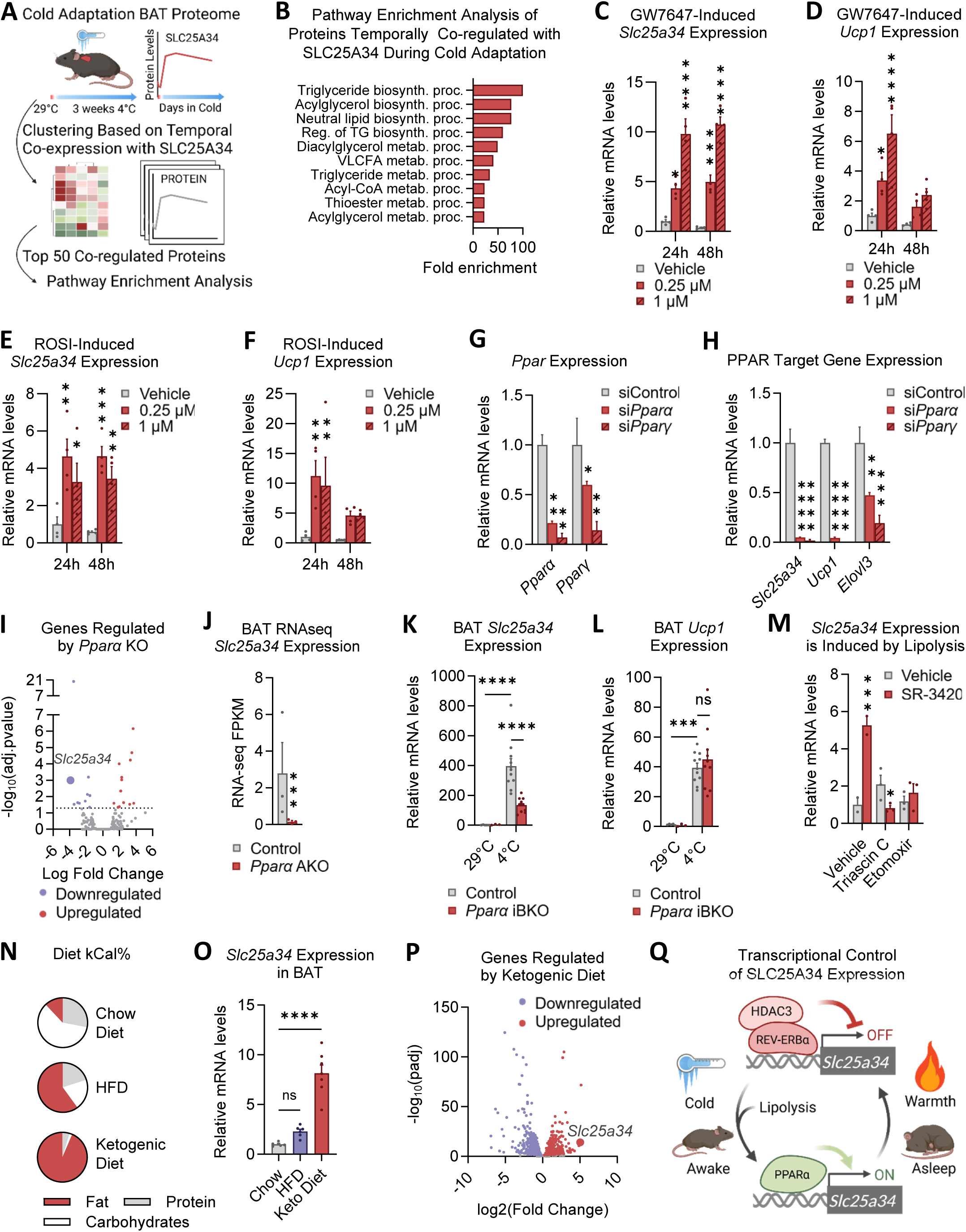
Cold-induced lipid metabolism activates PPARα to mediate temperature control of *Slc25a34*. **A**, schematic of the analysis design for identification of proteins temporally co-regulated with SLC25A34 in BAT during cold adaptation. **B**, Pathway enrichment of proteins temporally co-regulated with SLC25A34 in BAT during cold adaptation; VLCFA – very long chain fatty acids, TG - triglyceride. Regulation of *Slc25a34* and *Ucp1* mRNA by, **C-D**, PPARα agonist GW7647 or, **E-F**, PPARγ agonist rosiglitazone. **G-H**, gene expression in brown adipocytes following siRNA knockdown of *Pparα* and *Pparγ*. **I**, Volcano plot comparing BAT transcriptomes of control and *Pparα* AKO (targeted exon 5) mice acclimated to 22°C. **J**, *Slc25a34* mRNA expression in BAT of control and *Pparα* AKO (targeted exon 5) mice quantified in RNA-seq analysis. **K-L**, Gene expression in BAT from control and *Pparα* iBKO mice acclimated to 22°C and exposed to 29°C or 4°C for 3 days. **M**, *Slc25a34* mRNA in brown adipocytes following treatment with lipolytic activator, SR-3420, and the ACSL inhibitor, triascin C, or the CTP1 inhibitor, etomoxir. **N-O**, Diet composition and effects of diet challenges on BAT *Slc25a34* mRNA. **P**, Volcano plot depicting genes from BAT RNA-sequencing that are regulated by ketogenic diet. **Q**, schematic representation of how PPARα coordinates *Slc25a34* expression with REV-ERBα in BAT. For all panels, data are represented as mean ±SEM, p < 0.05 = *, p < 0.01 = **, p < 0.001 = ***, p < 0.0001 = ****, Two-way ANOVA with the Benjamini, Krieger, and Yekutieli two-stage linear step-up procedure for multiple comparisons correction (C-H, K-M), for (J) p-value was computed using Bioconductor software’s package and corrected for multiple testing using the Benjamini & Hochberg mode of the R function *p.adjust* to compute a false discovery rate (FDR), One-way ANOVA with the Benjamini, Krieger, and Yekutieli two-stage linear step-up procedure for multiple comparisons correction (O).

Therefore, we performed RNA sequencing of BAT from mice with adipose-specific *Pparα* knockout (*Pparα* AKO; targeting exon 5) and control littermates. Consistent with previously reported PPARγ compensation, only 10 BAT genes were significantly downregulated by *Pparα* deletion (**Fig. 2I, S2C, Table S5**). Strikingly, *Slc25a34* was the most downregulated of these 10 (**Fig. 2H, 2I, S2C**), revealing the transporter to be one of the few strong and uniquely selective PPARα targets. Neither *Ucp1* nor *Slc25a35* expression was affected by the *Pparα* deletion (**Fig. S2D, S2E**). Notably, *Pparγ* expression was unchanged in the *Pparα* knockout (**Fig. S2F**), indicating that it does not compensate for *Pparα* in the control of *Slc25a34* expression *in vivo*. We additionally confirmed the PPARα dependence of *Slc25a34* expression *in vivo* using a second adipose-specific KO model in which exon 4 was deleted (*36*) (**Fig. S2G-S2J**).

To address the effect of acute *Pparα* knockout on physiological BAT *Slc25a34* regulation, we used a tamoxifen-inducible *Pparα* knockout mouse model expressing CreERT2 under control of the *Ucp1* promoter (*Pparα* iBKO). *Pparα* iBKO and littermate control mice acclimated to room temperature (22°C) were exposed to thermoneutrality (29°C) or cold (4°C) for 3 days, concurrent with the tamoxifen-induced knockout. As confirmation of the knockout induction, *Pparα* expression was decreased in *Pparα* iBKO mice both at thermoneutrality and after cold exposure compared to controls (**Fig. S2K**). This acute ablation of *Pparα* in BAT significantly blunted the cold induction of *Slc25a34*, demonstrating the importance of PPARα for physiological temperature-sensitive regulation of *Slc25a34* (**Fig. 2K**). Conversely, cold-induced *Ucp1* expression, which is compensated by PPARγ, was unaffected by *Pparα* ablation (**Fig. 2L**).

The transcription-activating function of PPARs can be regulated by lipolytic products, which act as agonists (*37*, *38*). Given that lipid catabolism is robustly increased in BAT during cold exposure (*6*), we evaluated whether lipolysis influenced *Slc25a34* expression. Brown adipocytes were treated with SR-3420, a small molecule that directly enhances ATGL-dependent triglyceride lipolysis by disrupting the ABHD5–PLIN1 interaction (*39*, *40*). SR-3430 increased the expression of *Slc25a34* mRNA in the brown adipocytes (**Fig. 2M**). Furthermore, we completely prevented the lipolytic induction of *Slc25a34* (I) by blocking fatty acid activation, using the long-chain acyl-CoA synthetase (ACSL) inhibitor, triacsin C, or (II) by preventing transport of fatty acids into the mitochondria, using the carnitine palmitoyltransferase 1a (CPT-1a) inhibitor, etomoxir (**Fig. 2M**). These data indicate that endogenous lipid products from mitochondrial fatty acid oxidation activate PPARα-dependent control of *Slc25a34* in brown adipocytes.

Notably, exogenous dietary lipids are also known to activate PPARα and BAT thermogenic capacity, similarly to cold exposure (*41*). Therefore, we compared the effects of a 60% high fat diet (HFD) and a ketogenic diet with 93% fat content to chow diet (**Fig. 2N**) on the expression of the transporter in BAT of mice acclimated to room temperature (22°C). HFD modestly increased AAA, but the ketogenic diet, which has higher lipid content, caused a profound induction (**Fig. 2O**). Bulk RNA-sequencing further revealed that *Slc25a34* was the second most induced gene in the entire BAT transcriptome following ketogenic diet feeding (**Fig. 2P, Table S6**), suggesting a possible role for the transporter in this dietary paradigm and underscoring the impact of lipids on its expression. Collectively, our findings demonstrate that *Slc25a34* resides uniquely at the axis of circadian-mediated REV-ERBα-HDAC3 repression and cold-induced, lipid-liganded PPARα activation (**Fig. 2Q**).

### SLC25A34 controls mitochondrial respiration in thermogenic adipocytes

To investigate the role of the transporter in mitochondrial function, *Slc25a34* was acutely depleted in immortalized murine brown adipocytes. Three different siRNAs significantly decreased respiratory capacity, including basal and NE-stimulated respiration, and maximal respiratory capacity (**Fig. 3A, S3A, and S3B**). *Slc25a34* knockdown also reduced maximal respiratory capacity of the brown adipocytes not treated with NE (**Fig. 3B**). siRNA-mediated knockdown of *Slc25a34* also decreased NE-stimulated and maximal respiration in primary brown adipocytes (**Fig. 3C**). Conversely, knockdown of *Slc25a35* had little or no impact on NE-stimulated or maximal respiratory capacity (**Fig. 3D and S3C**), further highlighting a clear functional distinction between the two homologs.

**Figure 3.**
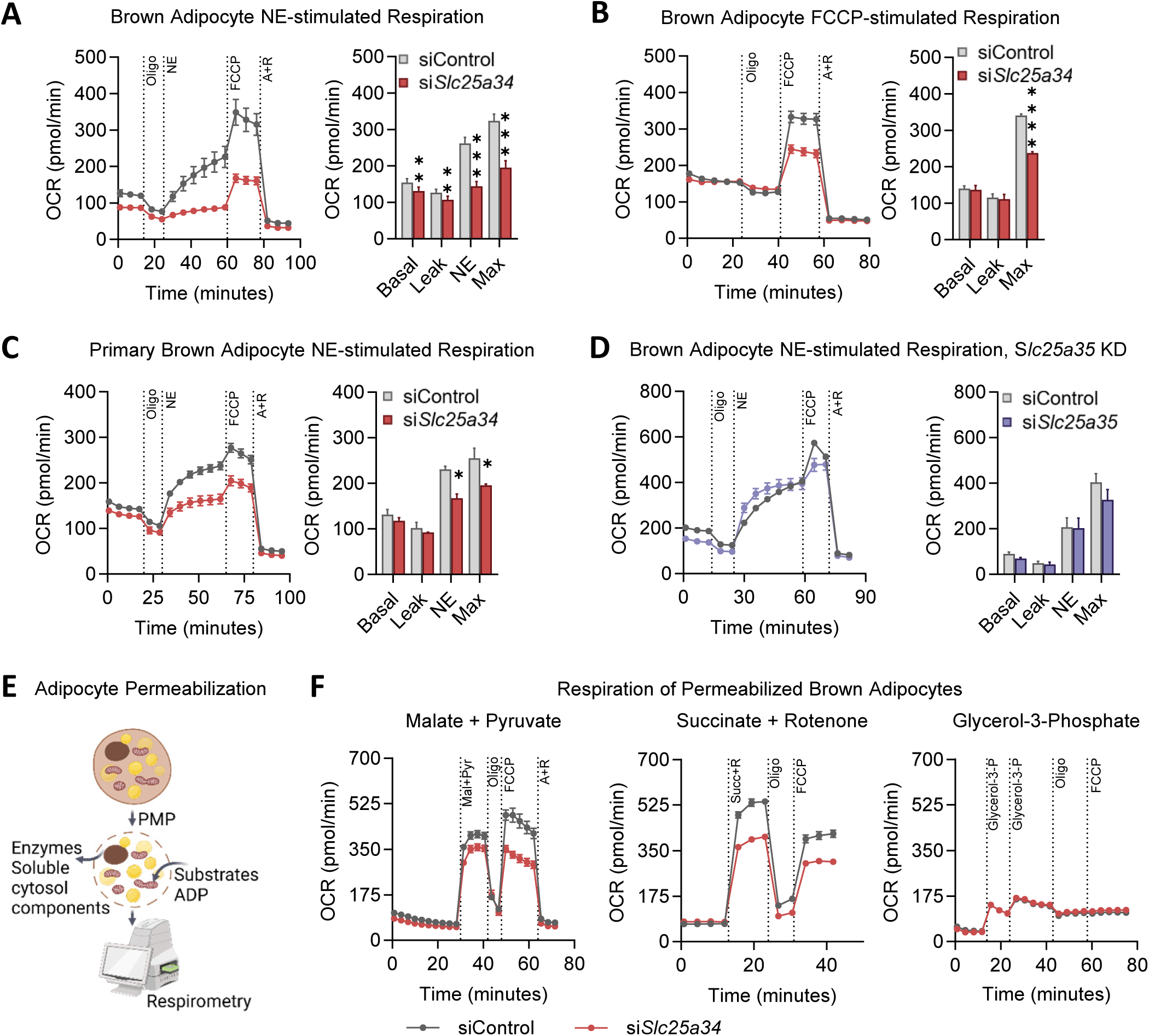
SLC25A34 controls mitochondrial respiratory capacity in thermogenic adipocytes. **A**, NE-induced and, **B**, maximal oxygen consumption rate (OCR) of brown adipocytes with control or siRNA-mediated *Slc25a34* knockdown; representative trace (left) and quantification of maximal OCR values reached during each run in 9 (A) and 3 (B) independent experiments (right). **C**, Oxygen consumption rate in primary brown adipocytes with control or siRNA-mediated *Slc25a34* knockdown; representative trace (left) and quantification of maximal OCR values reached during each run in 2 independent experiments (right). **D**, Oxygen consumption rate in primary brown adipocytes with control or siRNA-mediated *Slc25a35* knockdown; representative trace (left) and quantification of maximal values reached during each run in 4 independent experiments (right). **E**, Schematic of experimental design for assessing respiration in permeabilized brown adipocytes. **F**, respiration of permeabilized brown adipocytes following 4 days of siRNA-mediated *Slc25a34* knockdown and addition of specific respiratory substrates (from left to right): pyruvate + malate, succinate with addition of rotenone, and glycerol-3-phosphate. *Slc25a34* knockdown on panels A-C, F was performed with siRNA construct #1, measurements are made after 4 days of KD induction. For all panels, data are represented as mean ±SEM, p < 0.05 = *, p < 0.01 = **, p < 0.001 = ***, p < 0.0001 = ****, Paired multiple t-test with two-stage step-with the Benjamini, Krieger, and Yekutieli two-stage linear step-up procedure for multiple comparisons correction (A), Two-way ANOVA with the Benjamini, Krieger, and Yekutieli two-stage linear step-up procedure for multiple comparisons correction (B-D).

Mitochondrial metabolite carriers from the SLC25 family transport a diverse array of metabolites across the IMM, such as carboxylic acids, amino acids, fatty acids, nucleotides, and cofactors (*25*, *26*). Sequence analysis suggests that SLC25A34 is a carboxylic acid transporter (*25*, *26*). Given that several carboxylic acids serve as mitochondrial fuel, we measured the respiratory capacity of permeabilized brown adipocytes with control versus *Slc25a34* knockdown and supplemented with different substrates for respiratory complex I (malate, pyruvate, glutamate), complex II (succinate), or donating electrons into the Q-pool (octanoylcarnitine, palmitoylcarnitine, glycerol-3-phosphate) (**Fig. 3E**). We observed lower coupled respiration in the cells with the *Slc25a34* knockdown on nearly all the tested substrates (**Fig. 3F and S3D**), indicating that the respiratory deficiency was unlikely due to disrupted transport of any single substrate. Thus, our findings indicate that SLC25A34 is required for mitochondrial respiration of brown adipocytes.

### SLC25A34 promotes cytosolic acetyl-CoA production via mitochondrial import of oxaloacetate

To obtain unbiased insights into the potential function of SLC25A34, we profiled the metabolomes of brown adipocytes after four days of siRNA-mediated knockdown. Of the 56 differentially regulated metabolites (**Table S7**), the most significant change was the accumulation of N-acetylaspartate (**Fig. 4A, 4B**). N-acetylaspartate is synthesized in the mitochondria from oxaloacetate-derived aspartate and exported via an unknown mechanism to the cytosol where it plays a key role in thermogenic adipocytes by increasing the cytosolic acetyl-CoA pool through ACSS2 (*42*). Consistent with disrupted utilization of N-acetylaspartate, whole-cell acetyl-CoA levels, which largely reflect the cytosolic and nuclear concentrations (*43*), were significantly reduced following four days of *Slc25a34* knockdown (**Fig. 4C**), suggesting that *Slc25a34* plays a role in maintaining the acetyl-CoA pool in thermogenic adipocytes.

**Figure 4:**
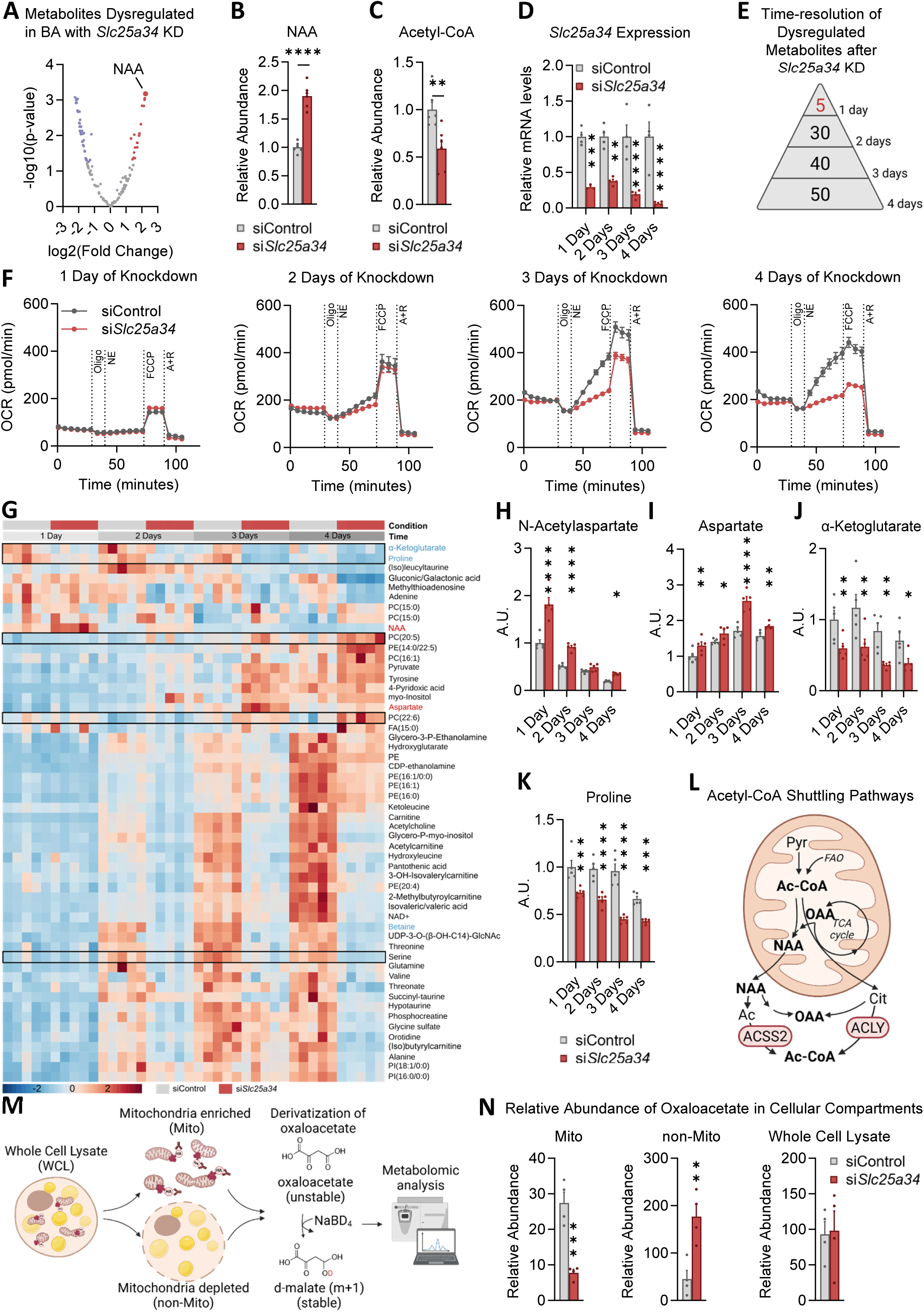
SLC25A34 promotes cytosolic acetyl-CoA production via import of oxaloacetate. **A**, Global differential metabolite changes between brown adipocytes with control or *Slc25a34* knockdown after 4 days of the knockdown induction. Levels of, **B**, N-acetylaspartate (NAA) and **C**, acetyl-CoA in brown adipocytes with control or *Slc25a34* knockdown after 4 days of the knockdown induction. **D**, *Slc25a34* expression, **E**, overview of metabolite changes, **F**, mitochondrial respiration, **G**, heatmap of metabolite changes, and **H-K**, specific metabolite levels from an *Slc25a34* siRNA-mediated knockdown time-course in brown adipocytes. **L**, Model showing how the generation of cytosolic acetyl-CoA by mitochondrial export of citrate (Cit) and NAA both lead to production of oxaloacetate (OAA) in the cytosol. **M**, Schematic for organelle-specific metabolomics and, **N**, abundance of OAA across cellular compartments. *Slc25a34* knockdown on all panels was performed with siRNA construct #1. For all panels, data are represented as mean ±SEM, p < 0.05 = *, p < 0.01 = **, p < 0.001 = ***, p < 0.0001 = ****, unpaired two-tailed Student’s t-test (B, C, N), Two-way ANOVA with the Benjamini, Krieger, and Yekutieli two-stage linear step-up procedure for multiple comparisons correction (D, H-K).

However, given the large number of metabolites that were changed four days after *Slc25a34* depletion, we next performed a time course with earlier knockdown time points to uncover precipitating metabolic changes. Yet even after just one day of knockdown, before any changes in cellular respiratory capacity manifested, N-acetylaspartate was still the most dysregulated among only five significantly altered metabolites (**Fig. 4D-H, Table S8, Table S9**), indicating it may be directly impacted by loss of SLC25A34. The other acute metabolite changes included increased aspartate (Asp), which was consistent with the higher N-acetylaspartate levels, and decreased α-ketoglutarate (α-KG), proline, and betaine (**Fig. 4I-4K, and S4A**). These changes further suggest that acute ablation of *Slc25a34* disrupts the malate-aspartate shuttle (MAS) and/or the TCA cycle. Importantly, the N-acetylaspartate-to-acetate route is not the only means of generating cytosolic acetyl-CoA in adipocytes. Another major pathway involves the mitochondrial export of citrate, which is converted to acetyl-CoA by ATP Citrate Lyase (ACLY) (*44*, *45*). Notably, both ACLY and ACSS2, the key enzymes in the two acetyl-CoA generating pathways, are dually regulated by REV-ERBα and cold temperature in BAT, similarly to SLC25A34 (**Fig. S4B**), providing a potential link between these pathways. Further underscoring this link, both the N-acetylaspartate and citrate routes of cytosolic acetyl-CoA production result in the generation of oxaloacetate in the cytosol (**Fig. 4L**), which is the proposed substrate of the yeast homologue of SLC25A34 (*28*, *46*). Therefore, we hypothesized that SLC25A34 could import oxaloacetate, generated in the cytosol from N-acetylaspartate and citrate, back into mitochondria to replenish the pool of TCA cycle intermediates.

To test whether the knockdown of the transporter affected oxaloacetate distribution in mammalian cells, we employed compartment-specific metabolomics using rapid HA-tag-based mitochondrial purification (*47*) from brown adipocytes with or without *Slc25a34* knockdown (**Fig. 4M**). Due to the labile nature of oxaloacetate, we chemically derivatized samples with sodium borohydride, converting oxaloacetate to a more stable compound, deuterated malate, for quantification. Total cellular oxaloacetate was unaffected by *Slc25a34* knockdown, however, cytosolic oxaloacetate accumulation increased significantly while the mitochondrial oxaloacetate pool was concurrently reduced (**Fig. 4N**). These findings are consistent with a role for SLC25A34 in facilitating oxaloacetate import back into mitochondria. Similar to our whole cell metabolomics, we again observed dysregulation of MAS metabolites, aspartate, glutamate, and malate (**Fig. S4C-S4E**), consistent with the established role of oxaloacetate in this circuit.

To explore the biochemical feasibility of oxaloacetate as a substrate for SLC25A34, we performed docking simulations of oxaloacetate with the human SLC25A34 in Autodock Vina using the SLC25A34 structure predicted by Alpha-fold. These simulations were performed with hydrated substrate and flexible residues in the conserved binding sites and matrix salt bridge network. We found that oxaloacetate could dock on SLC25A34 with robust affinity (−15.42 kcal/mol) in a binding pose in which it forms polar contacts between its carboxyl and carbonyl group and two arginine residues (R176 and R277), in the common substrate binding site of the transporter (**Fig. S4F**). Notably, this pose also permits the formation of polar contacts between the carbonyl and carboxyl groups of oxaloacetate and the four residues that make up the matrix salt bridge network (K31, D228, E28, and K128), which braces SLC25A34 in a matrix closed conformation. Docking to the conserved binding site in a manner that can disrupt the matrix salt bridge network is necessary for substrate transport by SLC25A carriers (*48*). Collectively, our findings support a model whereby mitochondrial import of oxaloacetate by SLC25A34 functionally links two major pathways for generating cytosolic acetyl-CoA.

### SLC25A34 couples interorganellar lipid cycling with transcriptional control of mitochondrial biogenesis

In adipocytes, one of the primary metabolic fates of cytosolic acetyl-CoA is to support *de novo* lipogenesis (DNL) (*49*). We assessed DNL through the incorporation of radiolabeled carbons from ^14^C-glucose into triglycerides after control or siRNA knockdown of *Slc25a34* (**Fig. 5A**). ^14^C-glucose was used because it radiolabels cytosolic acetyl-CoA, and, subsequently, triglycerides, through TCA-derived citrate via ACLY (*50*) and N-acetylaspartate via ACSS2 (*51*), the two rate-limiting enzymes we earlier found to be cold and circadian-regulated (**Fig. S4B**) similar to SLC25A34. Consistent with the acetyl-CoA measurements, loss of *Slc25a34* significantly diminished glucose-supported triglyceride production in brown adipocytes (**Fig. 5B**).

**Figure 5:**
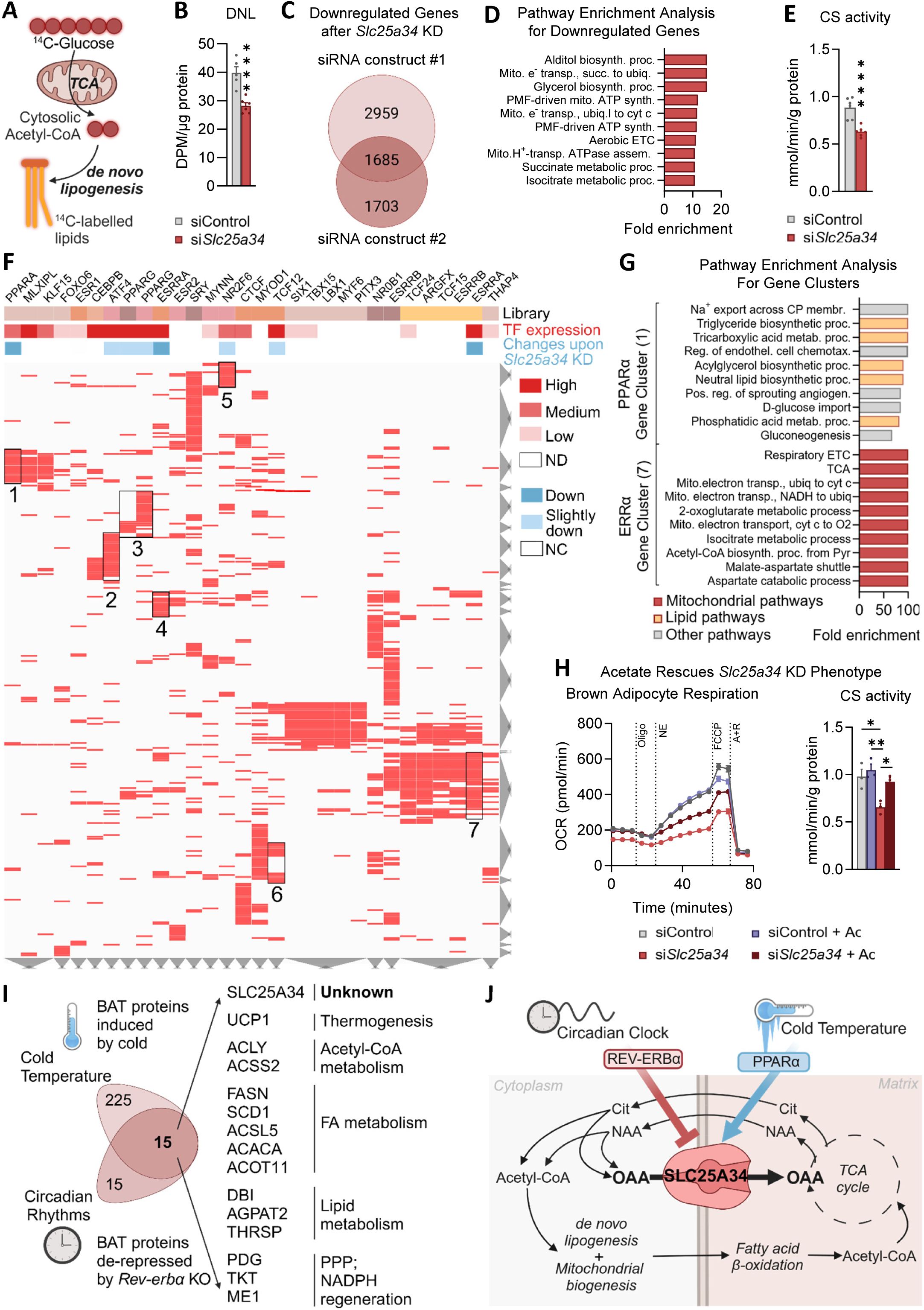
SLC25A34 couples interorganellar lipid cycling with transcriptional control of mitochondrial biogenesis. **A-B**, Schematic and assessment of radio-labeled glucose-derived *de novo* lipogenesis (DNL) in brown adipocytes with control or siRNA-mediated *Slc25a34* knockdown after 4 days of KD. **C**, Comparison of genes significantly downregulated following *Slc25a34* knockdown using two different siRNAs (Constructs #1 and #2). **D**, Pathway enrichment for the conserved gene signature downregulated by both siRNAs. **E**, Citrate synthase (CS) activity of brown adipocytes with control or *Slc25a34* knockdown after 4 days of KD. **F**, Promoter motif analysis of the genes downregulated by loss of *Slc25a34* for candidate transcription factors and, **G**, pathway enrichment of factor-specific gene clusters. **H**, Effect of acetate supplementation on respiration and mitochondrial content of brown adipocytes with siRNA-mediated *Slc25a34* knockdown; representative seahorse trace (left) and citrate synthase (CS) activity (right). *Slc25a34* knockdown was performed with siRNA Construct #1, measurements done after 4 days of knockdown induction. **I**, Classification of the protein cluster jointly regulated by cold and the circadian clock. **J**, Model of how the import of oxaloacetate (OAA) into mitochondria by SLC25A34 enables cytosolic acetyl-CoA production and intracellular lipid cycling. For all panels, data are represented as mean ±SEM, p < 0.05 = *, p < 0.001 = ***, p < 0.0001 = ****, unpaired two-tailed Student’s t-test (B, E), One-way ANOVA with the Benjamini, Krieger, and Yekutieli two-stage linear step-up procedure for multiple comparisons correction (H, right).

In addition to fueling DNL, cytosolic acetyl-CoA is also a critical mediator of transcriptional control of nuclear-encoded genes (*44*, *52*). To assess the potential impact of *Slc25a34*-dependent acetyl-CoA production on nuclear transcription, we globally profiled gene expression from brown adipocytes with or without knockdown of the transporter using two siRNA constructs (construct #1 and construct #2). When comparing the expression profiles of the two siRNAs, the shared changes revealed that 1,489 genes were increased and 1,685 genes were decreased (**Fig. S5A and 5C, Table S10**). Notably, despite causing a general reduction in acetyl-CoA levels, *Slc25a34* ablation specifically affected the expression of particular gene networks. Pathway enrichment analysis revealed that the upregulated genes were related to inflammatory and interferon pathways, underscoring the importance of SLC25A34 in maintaining cellular homeostasis (**Fig. S5B**). Whereas the downregulated genes were most significantly linked to mitochondrial biogenesis and respiratory capacity (**Fig. 5D**). Downregulation of these pathways was consistent with a decrease in mitochondrial mass, as assessed by citrate synthase activity (**Fig. 5E**). Taken together, our findings demonstrate that SLC25A34 impacts both mitochondrial function and transcriptional control of mitochondrial abundance.

As a transporter embedded in the IMM, SLC25A34 is unable to directly regulate nuclear genes. Therefore, to gain insight into the downstream transcriptional control, we interrogated the promoters of SLC25A34-regulated genes for transcription factor motifs. To refine our focus to the most relevant candidates, we further assessed how highly expressed each transcriptional regulator was in brown adipocytes and whether its expression was changed by *Slc25a34* knockdown. Among the genes that were increased following *Slc25a34* ablation, the promoters in the cluster representing inflammatory pathways were most enriched for STAT and IRF transcription factor motifs (**Fig. S5C, Table S11**). These factors were also highly basally expressed in brown adipocytes and were increased upon *Slc25a34* knockdown, suggesting that STATs and IRFs mediated induction of inflammatory pathways following loss of the transporter (**Fig. S5D**).

Analysis of the genes that were downregulated after loss of *Slc25a34* revealed seven clusters linked to transcription factors that satisfied the criteria for brown adipocyte expression and SLC25A34-dependency (**Fig. 5F, Table S11**). Pathway enrichment analysis found significant associations with three of the seven clusters. Clusters 1 and 3 represent targets of PPARα and PPARγ and were comprised of genes linked to lipid synthesis and catabolism (**Fig. 5G and S5E**). This SLC25A34-dependent regulation of PPAR lipid metabolism genes suggests a positive reinforcement mechanism, given our earlier finding that PPARα also drives *Slc25a34* expression. The genes in cluster 7 are related to mitochondrial biogenesis and respiratory function and were most strongly linked to ERRα control (**Fig. 5G**). Thus, loss of *Slc25a34* and the subsequent decrease in cytosolic acetyl-CoA appear to robustly impact nuclear transcription through specific transcription factor pathways.

We next investigated whether the role of SLC25A34 in supporting mitochondrial function at multiple levels depended on its ability to promote cytosolic acetyl-CoA production. Cytosolic acetyl-CoA can also be generated directly from exogenously added acetate via ACSS2, theoretically circumventing the requirement for mitochondrial export of N-acetylaspartate or citrate. Therefore, we hypothesized that supplementing cells with acetate might prevent the functional disruptions caused by *Slc25a34* knockdown (**Fig. S5F**). Indeed, we found that culturing brown adipocytes with acetate largely mitigated the respiratory defects caused by *Slc25a34* knockdown (**Fig. 5H, S5G-H**). Furthermore, acetate supplementation fully restored mitochondrial abundance, as indicated by citrate synthase activity (**Fig. 5H**). These findings underscore the importance of SLC25A34 in cytosolic acetyl-CoA production.

Collectively, our data demonstrate that SLC25A34 plays an essential role in both cytosolic DNL and mitochondrial respiration, thereby coordinating both anabolic and catabolic lipid processes across organelles. The importance of SLC25A34 in integrating interorganellar lipid metabolism is further highlighted when mapping the entire set of proteins that are co-regulated by cold and the circadian clock onto their respective metabolic pathways (**Fig. 5I**). Every one of these proteins is a part of a pathway that supports lipid synthesis, shuttling, or re-esterification as well as mitochondrial thermogenesis, and is functionally coupled between the cytosol and mitochondria by SLC25A34 (**Fig. S5I).** Taken together, our findings support a model whereby circadian and temperature-sensing programs in adipocytes leverage mitochondrial import of oxaloacetate via SLC25A34 to uniquely couple and accelerate the citrate and N-acetylaspartate pathways for generating cytosolic acetyl-CoA (**Fig. 5J**). This acetyl-CoA pool fuels DNL and transcriptional control of mitochondrial biogenesis, both of which serve to increase mitochondrial lipid oxidation and thermogenesis and, ultimately, complete an SLC25A34-dependent interorganellar circuit of lipid synthesis and breakdown. Notably, in humans, SLC25A34 appears to contribute to adipose function and whole-body energy homeostasis beyond thermogenic BAT depots. A meta-analysis of 23 clinical studies (https://adiposetissue.org/) (*53–76*) revealed that expression of *SLC25A34* in scWAT was negatively correlated with circulating C-reactive protein, TGs, and LDL cholesterol, HOMA-IR, BMI, waist circumference and waist-to-hip ratio, and positively correlated with circulating HDL cholesterol and the ratio of stimulated versus basal lipolysis (**Fig. 6**). Notably, there were no significant associations with *SLC25A34* expression in omental adipose tissue, suggesting that SLC25A34 may play a depot-specific beneficial role in promoting systemic metabolic health.

**Figure 6.**
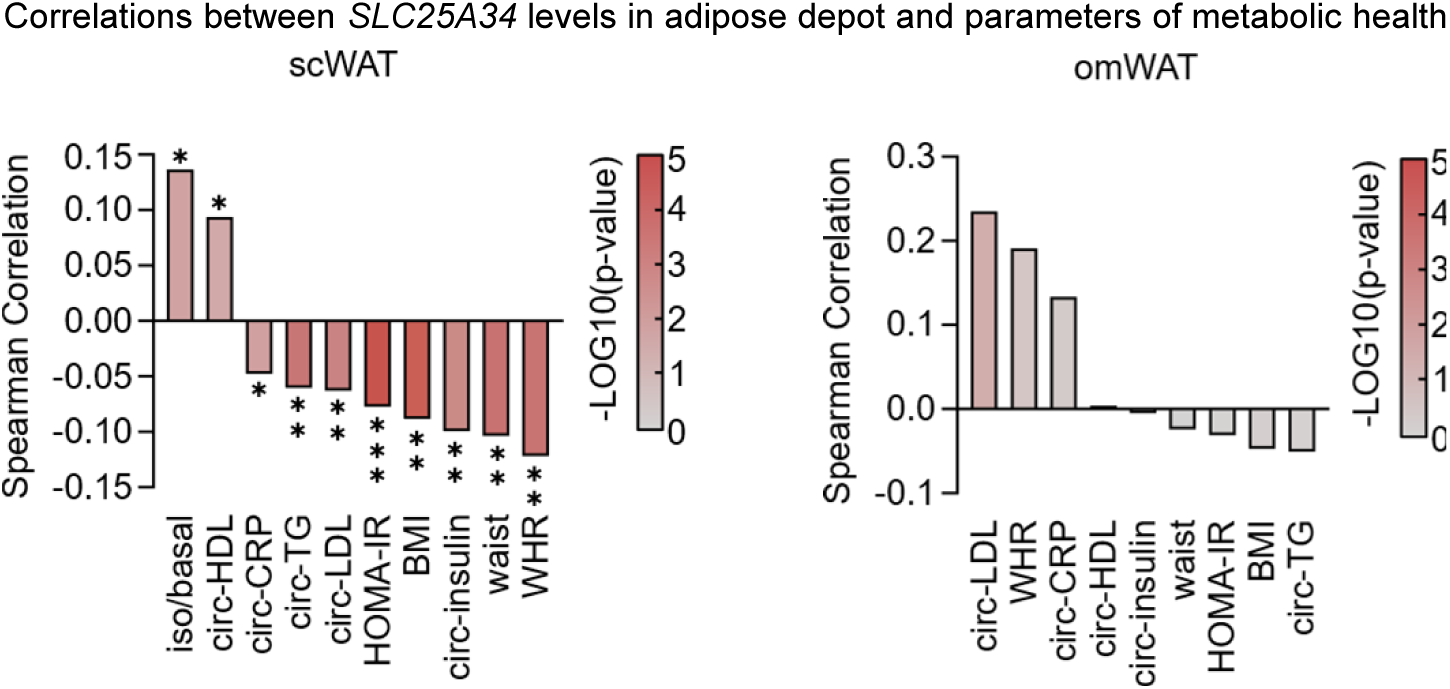
Correlations between WAT *SLC25A34* and parameters of metabolic health. Meta-analysis of 23 studies showing correlations between *SLC25A34 expression* in subcutaneous WAT (scWAT) and omental WAT (omWAT) and metabolic health parameters in human patients: ratio of stimulated versus basal lipolysis (iso/basal), circulating high-density lipoprotein (circ-HDL), circulating C-reactive protein (circ-CRP), circulating triglycerides (circ-TG), circulating low-density lipoprotein (circ-LDL), Homeostasis Model Assessment of Insulin Resistance (HOMA-IR), Body Mass Index (BMI), circulating insulin (circ-insulin), waist circumference (waist), and Waist-to-Hip Ratio (WHR).

## DISCUSSION

The circadian clock has been evolutionarily fine-tuned over hundreds of millions of years to anticipate recurring daily metabolic needs and coordinate tissue-specific homeostatic programs accordingly (*77*). To ensure maximal organismal fitness, this persistent rhythmic framework must be integrated with a means of rapidly responding to acute environmental and dietary changes. However, chronic modern-day perturbations, such as habitually eating during the sleeping period (*78*) or consuming diets high in fat content (*79*), extend beyond the protective capacity of these adaptations and can deleteriously impinge on circadian health. This is linked to the development of a range of diseases, including neurologic, psychiatric, cardiometabolic, and immune disorders (*80*). Therefore, delineating the molecular mechanisms that integrate circadian machinery with tissue-specific metabolic control is critical not only to understand evolutionary biology but also to identify key focal points for potential therapeutic intervention.

Adipose thermogenesis represents an ideal paradigm through which to study the integration of circadian, environmental, and dietary control of metabolism. Thermogenic functions are actively suppressed by the clock during sleep, then begin to increase just prior to awakening, reaching their peak during the waking period. The circadian rhythm of BAT activity plays a crucial role in regulating the daily fluctuations in systemic lipid homeostasis (*16*) and glucose consumption (*9*, *14*). Yet, the continuity of circadian rhythmicity must be coordinated with the rapid responsiveness of BAT activity to environmental cold exposure (*4*, *81*). Cold-induced BAT activity is also linked to the control of glucose and lipid metabolism (*17*). Finally, BAT activity is also shaped by dietary composition and caloric density, whereby lipid-rich diets induce thermogenic output (*82*). In the present work, we discover that the orphan mitochondrial metabolite carrier SLC25A34 represents a key regulatory node in BAT downstream of all three inputs by integrating circadian control, through REV-ERBα-HDAC3 repression, with cold temperature and dietary regulation, through lipid-mediated PPARα activation.

A hallmark of activated BAT is the simultaneous engagement of both mitochondrial fatty acid oxidation (FAO) and cytosolic DNL (*83*). This enables futile thermogenic cycling, through which FAO and glucose catabolism (*50*) feed the mitochondrial acetyl-CoA pool and support the TCA cycle. The mitochondrial TCA cycle-derived metabolites citrate and N-acetylaspartate are then exported to the cytosol, where they boost production of cytosolic acetyl-CoA, which serves as the priming substrate for DNL. Lipids from DNL can then be incorporated into triglycerides or transported back into the mitochondria to serve as FAO substrates and, thus, the cycle repeats. Notably, the critical step of generating cytosolic acetyl-CoA from either citrate or N-acetylaspartate produces an oxaloacetate in the cytosol whose metabolic fate has remained unresolved. We hypothesize that by directly shuttling oxaloacetate back into the mitochondria for re-entry into the TCA cycle, SLC25A34 would conceivably accelerate the futile lipid cycle.

Evidence for oxalacetate as a transport substrate comes from the closest deorphanized homolog of *Slc25a34*, *S. cerevisiae* mitochondrial carrier *OAC1*, which was shown to be able to directly transport oxalacetate (*46*). While our Alpha-fold docking studies support a model conducive to oxaloacetate as a substrate for mammalian SLC25A34, *in vitro* studies using recombinantly expressed SLC25A34 in reconstituted liposomes will be needed to unequivocally demonstrate oxaloacetate transport. Nevertheless, our collective findings show that SLC25A34 is functionally required for both DNL and mitochondrial oxidation and, thus, appears to represent a novel mechanistic link in thermogenic lipid cycling.

In addition to promoting DNL and linking anabolic and catabolic lipid pathways through macronutrient exchange across the IMM, SLC25A34 also influences the transcription of nuclear-encoded genes related to lipid metabolism and mitochondrial biogenesis. The question remains as to whether this transcriptional control is directly downstream of cytosolic acetyl-CoA production, which is an enzymatic substrate for histone acetyltransferases, or if DNL is involved in producing a signaling molecule that regulates gene expression. Support for the latter mechanism comes from the specificity of the transcriptional response we observed following *Slc25a34* depletion and that the motif enrichment suggested the involvement of PPARα and PPARγ, which are liganded by products of lipid metabolism. Such a paradigm would confer the circadian clock, diet, and temperature-sensing pathways with the ability to simultaneously coordinate lipid oxidation and mitochondrial abundance.

Given its central importance to cellular metabolism, signaling, and transcriptional control, the ability to defend and maintain cytosolic acetyl-CoA levels is critical for the physiological function of numerous cell types across many tissues (*52*) and is exploited for the survival of certain types of cancer (*84*). Underscoring the essential roles of cytosolic acetyl-CoA, there are multiple pathways that feed into and support this metabolite pool (*52*). Our findings reveal that SLC25A34 functionally links two of the most prominent of these pathways, mediated by citrate and N-acetylaspartate mitochondrial export.

While the present study focuses on adipose thermogenesis, REV-ERBα and PPARα, the primary regulators of *Slc25a34* expression, have a multitude of key roles in numerous other tissues (*85–87*). Therefore, the impact of SLC25A34 likely extends beyond adipocytes. For example, mitochondrial transport of oxaloacetate has been observed in the heart, kidney, and liver (*88–93*). In fact, heart is the tissue with the highest expression of *Slc25a34*, further suggesting a role in cardiac biology. There is also a potential contribution to neuronal health given that N-acetylaspartate, which we found to be one of the BAT metabolites most strongly impacted by loss *Slc25a34*, is most canonically known for its role in the CNS. It is the second most abundant metabolite in the brain and supports myelination and neuronal integrity (*94*). However, ultimately, if and to what extent the metabolic signature of SLC25A34 in thermogenic adipocytes is conserved across these functionally distinct cell and tissue types remains to be determined.

## Acknowledgments

We thank the members of the Gerhart-Hines lab for discussions. We thank Katharina Stohlmann and Olivia Sveidahl Johansen for the technical assistance with mouse and cell studies. We acknowledge Thomas Gade Koefoed, Rikke Kammersgaard Nysted, Lars Roed Ingerslev, Mie Mechta, and the Single-Cell Omics Platform at the Novo Nordisk Foundation Center for Basic Metabolic Research for technical and computational expertise and support. We acknowledge Charlotte Helene Plass Hartung, Mansa Nair Kihara, and the Metabolomics Platform at the Novo Nordisk Center for Basic Metabolic Research, University of Copenhagen, for mass spectrometry-based metabolomics analyses. Figures 1A, 1K, 2A, 2Q, 3E, 4L, 4M, 5A, 5C, 5I, 5J, S5A, S5F, S5G, S5I were created with BioRender.com.

## Author contributions

Conceptualization: ZGH, IK, and MAL

Investigation and data analysis: IK, ALB, SAJT, MFH, LKM, JFH, MSI, HJR, ZK, YD, SC, YS, DN, FS, FF, LAM, DT, TM, EGS, HE, CKK, RLM, GJM, ASH, MJE, ZAK, MMF, MPJ, M van W, HM, EZ, RWMcG, KP, MKM, AP, PC, JGG, PS, RHH, JBH, SPG, TWS, MPG, TDHJ, RES, PJW, ERSK, TM, JTT, SM, BE, LK, NJF, MAL, ZG-H

Visualization: IK, ALB, YD, ZGH Writing – original draft: IK

Writing – review & editing: ZGH, IK, ALB, and SAJT (first draft). All authors read and edited the manuscript.

**Competing interests**

TWS and ZGH work or have worked, in some capacity, for Embark Biotech ApS, a company developing therapeutics for the treatment of diabetes and obesity. All other authors declare no competing interests associated with this manuscript.

## Data and materials availability

All data are available in the main text or the supplementary materials. RNA-seq and ChIP-seq data have been deposited to the NCBI Gene Expression Omnibus under accession number <u>GSEXXXXXX</u>.

## Materials and Methods

### ANIMAL MODELS

Wild type (C57Bl/6JBomTac) mouse studies (Tissue panel (1B, S1A), Ketogenic diet study (Fig. 2N, 2O), cold acclimation study (Fig. 1C, 1D, 1E, S1B), and mice used for primary adipocyte culture) were performed with approved protocols from The Danish Animal Experiments Inspectorate (permit number: 2018-15-0201-01441) and the University of Copenhagen (project number: P18-312 and P19-374). Mice were housed in an enriched environment with ad libitum access to standard chow diet (Altromin, 1310) and water. Light in the facility was set to a 12 h light/dark cycle (light: 6 AM-6 PM and dark: 6 PM-6 AM) at 22±2°C. Studies were performed in male and female mice between 8 and 20 weeks of age. Mice were housed at room temperature (RT, 22±2°C) unless otherwise stated. Housing temperatures used in the studies: thermoneutrality - 29°C, room temperature – 22±2°C, cold – 4-5°C.

Details of the housing conditions and studies performed on *Rev-erbα* WKO, *Rev-erbα/β* dBKO, *Hdac3* AKO, *Pparα* AKO and *Pparα* iBKO mice are specified below.

### Tissue panel

The tissue panel experiment was performed as described in (*95*). Briefly, 20 weeks old C57Bl/6JBomTac (Taconic Biosciences) mice were acclimatized to thermoneutrality for 3 weeks prior to the 24 hours cold challenge at 4°C. Tissues were harvested at the end of cold challenge between 9 AM (ZT3) and 1PM (ZT7).

### Cold acclimation study

This study is published in (*24*). For cold exposure experiments, 12- to 16-week-old mice were individually housed and placed into climate-controlled rodent incubators (Memmert HPP750Life) set to 29°C to acclimate to thermoneutrality for two weeks. Cold exposure mice were then moved to an incubator set to 4°C and kept there until dissection. Tissues were harvested between 9 AM (ZT3) and 1PM (ZT7). BAT was used for proteomics analysis and gene expression analysis with qPCR.

### HFD and Ketogenic Diet study

(Fig. 2M, 2N)

Adult (8 weeks) male C57BL/6J mice were purchased from Janvier Breeding Centre (France), and either placed on a standard chow diet (Brogaarden, Denmark) or a 60% high fat diet (HFD) (Reasearch Diets) for 24 weeks. The mice were group-housed with up to 8 mice per cage at 22°C on a 12:12 h light-dark cycle. The mice had free access to food and water. Before the ketogenic diet study all mice were singly housed with 2 weeks for acclimatization. For the consecutive three weeks the mice from the chow diet either continued receiving chow diet (5 mice) or were given ketogenic diet (BioServ # F3666, NJ, USA), fresh ketogenic diet was given to the mice every second day. HFD group (6 mice) continued receiving HFD for the 3 weeks. All mice were fed ad libitum and therefore the food intake was not isocaloric. After 27 weeks since the beginning of the diet intervention the mice were sacrificed, and BAT was used to measure gene expression.

### Ketogenic diet study

(Fig. 2O)

Adult (14-weeks old) C57Bl/6N male mice acclimated to 22°C and fed with standard chow diet (Brogaarden, Denmark) were singly housed for 2 weeks prior to the experiment. For 3 weeks, the mice were given either a chow diet (4 mice) or a ketogenic diet from BioServ (F3666, NJ, USA) (4 mice) ad libitum. Fresh ketogenic diet was given to the mice every second day. The food intake between the two groups was not isocaloric. After three weeks of diet intervention, the mice were sacrificed and tissues harvested.

#### Rev-erbα WKO

Studies with the *Rev-erbα* whole body knockout mice (*Rev-erbα* WKO) were performed with an approved protocol from the University of Pennsylvania Perelman School of Medicine Institutional Animal Care and Use Committee as described in (*9*). The *Rev-erbα* WKO mice were obtained from B. Vennström and backcrossed seven or more generations with C57Bl/6 mice. Mice were housed on a 12:12-h light-dark cycle (lights on at 7 AM, lights off at 7 PM). Gene expression and protein analysis were carried out on 12-16 week old male *Rev-erbα* WKO mice and WT littermates.

For the circadian gene expression experiment (Fig. 1F) mice were singly housed at 29°C prior to the experiment. Tissues were harvested at ZT-1, ZT3, ZT7, ZT11, ZT15, ZT19, ZT23, ZT27 (2 to 4 animals per group per time point). For the proteomics study (Fig. 1G), mice were acclimated to thermoneutrality at 29°C, and tissues were harvested at 5 PM (ZT 10) (**Table S2**). The cold exposure study (Fig. 1H), was performed in climate-controlled rodent incubators. All WT and *Rev-erbα* WKO mice used in the studies were first placed in individual cages with access to food and water and allowed to acclimate to 29°C for 2 weeks prior to cold challenge. For the cold challenge mice were exposed to cold at 4°C at 11 AM (ZT 4) for 1.5 hours and tissues were harvested at ZT5.5.

#### Rev-erbα/β dBKO

All animal work was approved by the University of Pennsylvania Perelman School of Medicine Institutional Animal Care and Use Committee (IACUC protocol number 804747 issued to Mitchell Lazar). Male *Rev-erbα^fl/fl^/Rev-erbβ^fl/fl^* (*96*) mice were maintained on a C57Bl6/J background (Jackson Labs, Stock 008661). To produce *Rev-erbα/β* dBKO, *Rev-erb*α^fl/fl^/*Rev-erb*β^fl/fl^ mice were bred to Ucp1-Cre mice maintained on a C57BL/6J background (Jackson Labs Technologies, Inc., B6.FVB-Tg[Ucp1-Cre]1Evdr/J, Stock 024670) (*97*). Mice were bred and maintained at 20–22 °C on a 12:12 h light:dark cycle with *ad libitum* access to food and water unless otherwise noted. The normal chow diet (NCD, LabDiet, 5010) consisted of 12.7% kcal fat, 28.7% kcal protein, and 58.5% kcal carbohydrates. Gene expression was carried out on 11-16 week old male *Rev-erbαev-erbβ* dBKO mice and WT littermates. Cold exposure experiments were performed in climate-controlled rodent incubators set to 29°C and 4°C. Mice from the thermoneutrality acclimated group (5 WT and 5 KO mice) were housed at 29°C for 2 weeks and the tissues were harvested at 5 PM (ZT10). Mice from the cold challenge group (5 WT and 6 KO mice) were singly housed at 29°C for 4 days and acutely exposed to 4°C cold in pre-chilled cages at 4 °C with bedding and free access to standard chow, without a cotton nestlet or access to water, and cage lid partly open. Cold challenge lasted for 6 hours from 11 AM (ZT 4) to 5 PM (ZT 10). Tissues were harvested at 5 PM (ZT10) and BAT was used for RNA sequencing (**Table S3**).

#### Hdac3 AKO

Studies with *Hdac3* AKO mice were performed according to ethical regulations and protocols approved by the Institutional Animal Care and Use Committee (IACUC) of the Perelman School of Medicine at the University of Pennsylvania and the Children’s Hospital of Philadelphia as described in (*98*). Mice were group housed with enrichment in a temperature and humidity controlled, specific-pathogen free animal facility at 22°C under a 12:12-h light-dark cycle with free access to standard chow (LabDiet, 5010) and water. To generate the mouse strain with the adipose-specific *Hdac3* knockout (*Hdac3* AKO), *Hdac3*^f/f^ mice maintained on a C57BL/6 background were bred to B6;FVB-Tg(Adipoq-Cre)1Evdr/J mice form Jackson laboratory (stock No: 010803) (*98*). All experiments used male age-matched littermates group housed at 4–5 animals per cage. Gene expression studies were carried out on 10–12-week-old male mice at ZT 10. Cold-exposure experiments were performed in climate-controlled rodent incubators (Powers Scientific) maintained at 22°C or 4-5°C, with interior temperature monitoring by digital thermometers. For the cold exposure experiment, mice were allowed to acclimate to 22°C for two weeks. Cold exposure mice were placed in pre-chilled cages at 4–5 °C with bedding, a cotton nestlet, free access to standard chow and water, and the cage lid partly open at 5PM (ZT10). Mice were sacrificed and tissues harvested after 3 hours of cold exposure. Animal numbers were 4 WT and 3 KO in the room temperature group, 5 WT and 6 KO in the cold group).

#### *Pparα* AKO (exon 5 targeted)

All experiments with Pparα AKO mice with targeted exon 5 were performed according to procedures approved by the University of Pennsylvania’s Institutional Animal Care and Use Committee as described in (*33*). Male wild-type C57BL/6J (stock 000664) mice, as well as F1 intercrossed progeny with 129S1/SvlmJ (B6129SF1/J, stock 101043), were purchased from Jackson Laboratory. *Pparα*^flox/flox^ mice were kindly provided by F.J. Gonzales (National Institutes of Health, Bethesda) and have been described previously (*99*). To generate *Pparα* AKO mice, we crossed Adiponectin-Cre mice with *Pparα*^flox/flox^ mice. All mice were housed, 5 per cage, in a temperature-controlled, specific pathogen–free facility with a 12-hour light / 12-hour dark cycle and *ad libitum* access to water and food at room temperature (22±2°C). 12 weeks old male mice were sacrificed and BAT was used for RNA sequencing (3 WT and 3 KO mice).

#### *Pparα* AKO (exon 4 targeted)

All experimental procedures and protocols for *Ppara* AKO (exon 4) mice complied with the National Institutes of Health Guide for the Care and Use of Laboratory Animals and were approved by the Institutional Animal Care and Use Committee of the University of Mississippi Medical Center. The *Ppara* AKO (exon 4) mice were previously generated and described in (*36*). In short, homozygous *Ppara*^flox/flox^ (from Dr. Walter Wahli, first published in (*100*)) and heterozygous *Adipoq*-Cre mice were crossed to create *Ppara* AKO (exon 4) mice and *Ppara*^flox/flox^ littermate controls. Male and female mice were housed under standard conditions at room temperature with full access to water until 8 weeks of age. Mice were then switched to a normal chow (NCD) consisting of a 17% fat diet (Teklad 22/5 rodent diet, #860, Harland Laboratories, Inc., Indianapolis, IN, USA) for 30 weeks and had access to food and water *ad libitum*. After 30 weeks, mice were euthanized via overdose of isoflurane anesthesia, and tissues were immediately collected and frozen in liquid nitrogen. Tissue samples were stored at −80 °C until use.

#### Pparα iBKO

The study with *Pparα* iBKO mice were approved by the Danish Animal Experiment Inspectorate (approval #2018-15-0201-01459) and complied with the ARRIVE guidelines. Male mice of 7-8 weeks old were housed at 22°C, under a 12:12-h light-dark cycle, with free access to food and drinking water. To generate mice with brown adipose-specific *Pparα* knockout (*Pparα* iBKO), *Pparα*^f/f^ mice (kindly provided by F.J. Gonzales, National Institutes of Health, Bethesda) (*99*) were crossed with the tamoxifen-inducible B6-Tg(Ucp1-cre/ERT2)426Biat mouse line (kindly provided by Prof. Christian Wolfrum) (*101*). For the cold exposure experiment, RT acclimated mice were given 2 mg of tamoxifen (Sigma-Aldrich T5648) in 100 μl of corn oil (Sigma-Aldrich C8267) per mouse by oral gavage once per day for 2 consecutive days. Then the mice were spilt into 2 groups and transferred to 29°C or 4°C for 3 days, after which the tissue was harvested. One more tamoxifen gavage was given to all mice on the first day of the temperature challenge. Control mice were *Pparα*^f/f^ without Ucp1-Cre/ERT2 and were given an identical dosage of tamoxifen.

### CELL MODELS

#### Immortalized brown adipocyte culture

Murine brown preadipocytes immortalized with SV40 large T antigen (referred to as BMC) were kindly provided by Associate Professor Patrick Seale (*102*). Preadipocytes were cultured in basal DMEM (Gibco, 31966) containing 10% FBS (Sigma-Aldrich, F7524) and 1% penicillin/streptomycin. Cells were passaged when they reached 60-70% of confluence. Media was changed every 2nd day. On the day of 100% confluency, cell differentiation was induced by supplementing of standard media with the differentiation cocktail (insulin (20 nM) (Sigma-Aldrich, I9278), dexamethasone (1 mM) (Sigma-Aldrich, D1756), rosiglitazone (0.5 mM) (Cayman Chemicals, 71740), T3 (1 nM) (Sigma-Aldrich, T6397), and 3-isobutyl-1-methylxanthine (IBMX) (250 μM) (Sigma-Aldrich, I5879)). On day 2 of differentiation, media was switched to maintenance media containing insulin and T3. Maintenance media was changed every 2nd day. Cells were harvested or assayed on day 7 of differentiation unless stated otherwise in the figure legends. The cells were maintained at 37°C in a humidified atmosphere with 5% CO2.

#### Immortalized brown adipocyte culture (Figure 2L)

Murine brown preadipocytes immortalized with SV40 large T antigen (BMC) were kindly provided by Associate Professor Patrick Seale (*102*). Preadipocytes were cultured in basal DMEM (Sigma-Aldrich, D6429) containing 10% FBS (Sigma-Aldrich, F7524) and 1% penicillin/streptomycin. Cells were passaged when they reached 60-70% of confluence. Media was changed every 2nd day. On the day of 100% confluency, cell differentiation was induced by supplementing of standard media with the differentiation cocktail (insulin (20 nM) (Sigma-Aldrich, I9278), dexamethasone (0.5 mM) (Sigma-Aldrich, D4902), T3 (1 nM) (Sigma-Aldrich, T6397), and 3-isobutyl-1-methylxanthine (IBMX) (0.5 mM) (Sigma-Aldrich, I5879)). On day 2 of differentiation, media was switched to maintenance media containing insulin and T3. Maintenance media was changed every 2^nd^ day. Cells were harvested or assayed on day 7 of differentiation. The cells were maintained at 37°C in a humidified atmosphere with 5% CO2.

#### Primary brown adipocyte culture

Primary brown preadipocytes were isolated from wild type C57BL/6J male and female mice as described in (*95*). Male and female tissues were pooled for the generation of primary cell culture. BATs were harvested from 6-8 weeks old mice and put into ice-cold DMEM (Gibco, 31966) media. Tissues were minced and subsequently digested with DMEM (Gibco, 31966) supplemented with 2% BSA (Sigma-Aldrich, A7030) and 0.2% collagenase type I (Worthington Biochemical Corp., LS004197) at 37°C for 40 min. The digests were centrifuged (5 min, 400 g) to obtain the stromal vascular fraction (SVF) and pellets were resuspended in DMEM with 10% FBS (Sigma-Aldrich, F7524) and 1% penicillin/streptomycin. The cell suspensions were passed through a 70 μm strainer and distributed onto the plate-format of interest. On day 1 after cell harvest, the isolated preadipocytes were rinsed with DMEM containing 10% FBS and 1% penicillin/streptomycin and thereafter, media was changed every 2^nd^ day. The cells were passaged up to one time before differentiation. On the day of 100% confluency, cell differentiation was induced by supplementing of standard media with the differentiation cocktail (insulin (86 nM) (Sigma-Aldrich, I9278), dexamethasone (0.1 μM) (Sigma-Aldrich, D4902), rosiglitazone (0.5 mM) (Cayman Chemicals, 71740), T3 (1 nM) (Sigma-Aldrich, T6397), and 3-isobutyl-1-methylxanthine (IBMX) (250 μM) (Sigma-Aldrich, I5879)). On day 2 of differentiation, media was switched to maintenance media containing insulin and T3. Maintenance media was changed every 2^nd^ day. Cells were harvested or assayed on day 7 of differentiation. The cells were maintained at 37°C in a humidified atmosphere with 5% CO2.

#### siRNA mediated gene knockdown

Primary and immortalized murine brown preadipocytes were differentiated as stated above. On day 3 of differentiation, cells were reverse transfected according to (*103*). In brief, siRNA targeting the gene of interest or relevant control siRNA were diluted in Opti-MEM (Gibco, 51985) to a final concentration of 300 nM. RNAiMAX (Invitrogen, 13778-150) was added to a final concentration of 30 μl/ml. The siRNA mix was added to the bottom of the wells in the plate-format of interest and allowed to incubate at RT for 30 min. In the meantime, cells were trypsinized, counted, and resuspended in culture media. Finally, the cell suspension was distributed on top of the siRNA mix to a final siRNA concentration of 50 nM. Media was changed 2 days after transfection and cells were harvested or assayed on day 7 of the differentiation unless otherwise is stated in the figure legends. siRNA constructs used in the study are listed in **Table S13**.

#### Acetate rescue study

For the acetate rescue study immortalized brown adipocytes were grown on 10 cm cell culture dishes. On the day of confluency, the culture media was exchanged to differentiation media for control dish and to differentiation media containing 10 mM sodium acetate for the acetate treated dish. Differentiation protocol was standard for both groups. On day 3 of differentiation siRNA transfection was performed as described abouve. Acetate-treated group always received culture media freshly supplemented with 10 mM sodium acetate during transfection and regular media changes. Seahorse respirometry and was performed on day 7 in seahorse media containing standard respiratory substrates, without sodium acetate supplementation. Cells were harvested for CS activity assay on day 7.

### PROTEOMICS

Proteomics was performed exactly as described in (*24*). BAT proteomics data from the cold adaptation study is published and provided as Supplementary Material Data S1 in (*24*). BAT proteomics from *Rev-erbα* WKO mice is provided in Table S1.

Brown adipose tissues were mechanically lysed with a homogenizer with 2 mL SDS lysis buffer containing 2.0 % SDS w/v, 150 mM NaCl, PhosStop (Roche, Madison, WI) phosphatase inhibitors, EDTA free protease inhibitor cocktail (Promega, Madison, WI) and 50 mM HEPES, pH 8.5. Lysates were reduced with 5 mM DTT and cysteine residues were alkylated at room temperature with iodoacetamide (14 mM) in the dark as previously described (*104*). Protein content was purified by methanol/chloroform extraction. Protein disks were resuspended in 8 M Urea containing 50 mM HEPES (pH 8.5) and concentrations were measured by BCA assay prior to protease digestion. 1 mg of protein lysates were diluted to 4 M urea and digested with LysC (Wako, Japan) in a 1/100 enzyme/protein ratio overnight. Protein extracts were diluted further to a 1.0 M urea concentration and trypsin (Promega, Madison, WI) was added to a final 1/200 enzyme/protein ratio for 6 hours at 37°C. Digests were acidified with 20 uL of 20% formic acid (FA) to a pH ∼ 2 and subjected to C18 solid-phase extraction (SPE) (Sep-Pak, Waters, Milford, MA).

Isobaric labeling of peptides was performed using 10-plex tandem-mass tag (TMT) reagents (Thermo Fisher Scientific, Rockford, IL). TMT reagents (0.8 mg) were dissolved in 42μl dry acetonitrile (ACN) and 10 μl was added to 100 μg of peptides dissolved in 100 μl of 200mM EPPS, pH 8.0. After 1hr (RT), the reaction was quenched by adding 4 μl of 5% hydroxylamine. Labeled peptides were combined, acidified with FA (pH ∼2) and diluted to a final ∼5% ACN concentration prior to C18 SPE on Sep-Pak cartridges (50 mg).

Basic pH reversed-phase HPLC (bpHrp) TMT labeled peptides were subjected to orthogonal bpHrp fractionation. Labeled peptides were solubilized in buffer A (5% ACN 10 mM ammonium bicarbonate, pH 8.0) and separated by an Agilent 300 Extend C18 column (5 μm particles, 4.6 mm ID and 220 mm in length). Using an Agilent 1100 binary pump equipped with a degasser and a photodiode array (PDA) detector (Thermo Scientific, San Jose, CA), a 50 min linear gradient from 12% to 45% acetonitrile in 10 mM ammonium bicarbonate pH 8 (flow rate of 0.8 mL/min) separated the peptide mixtures into a total of 96 fractions. 96 Fractions were consolidation into 12 samples, acidified with 10 μl of 20% formic acid and vacuum dried. Each sample was re-dissolved in 5% formic acid, desalted via StageTips, dried via vacuum centrifugation, and reconstituted for LC-MS/MS analysis.

Mass spectrometry analysis: All bpHrp fractions were subjected to LC-MS/MS analyses onto a LTQ Orbitrap Velos Fusion (Thermo Scientific San José, CA) instrument equipped with a Famos autosampler (LC Packings, Sunnyvale, CA) and an Agilent 1100 binary HPLC pump (Agilent Technologies, Santa Clara, CA). Peptides were separated onto a 100 μm I.D. microcapillary column packed first with approximately 1 cm of Magic C4 resin (5μm, 100 Å, Michrom Bioresources, Auburn, CA) followed by ∼25 cm of Maccel C18AQ resin (1.8 μm, 200 Å, Nest Group, Southborough, MA). Peptides were separated by applying a gradient from 5 to 35% ACN in 0.5% FA over 180 min at ∼200 nl/min. Electrospray ionization implemented through applying a voltage of 1.86 kV using an inert gold electrode via a PEEK junction at the end of the microcapillary column. The LTQ Orbitrap Velos Fusion was operated in data-dependent manner for the MS methods. The MS survey scan was performed in the Orbitrap in the range of 400-1300 m/z at a resolution of 3x104, followed by the selection of the ten most intense ions (TOP 10) for CID-MS2 fragmentation in the ion trap using a precursor isolation width window of 2 m/z, AGC setting of 2000, and a maximum ion accumulation of 150 ms. Normalized collision energy was set to 35% and an activation time of 20ms. Ions within a 10 ppm m/z window around ions selected for MS2 were excluded from further selection for fragmentation for 120s. Directly following each MS2 event, 6-10 of most intense fragment ion in an m/z range between 110-160% of the precursor m/z was selected for HCD-MS3 (*105*, *106*). The fragment ion isolation width was set to 4 m/z, AGC was set to 20,000, the maximum ion time was 250ms, normalized collision energy was set to 60% and an activation time of 50ms for each MS3 scan. For all MS3 scans, Orbitrap resolving power was set to 30,000 (@ 400 m/z).

Mass spectrometry analysis: Data processing. A compilation of in-house software was used to convert mass spectrometric data (Raw file) to a mzXML format, as well as to correct monoisotopic m/z measurements and erroneous peptide charge state assignments. Assignment of MS/MS spectra was performed using the Sequest algorithm by searching the data against a protein sequence database including all entries the Mouse Uniprot database (download date June, 2014) containing known contaminants such as human keratins and its reverse decoy components (*107*). Sequest searches were performed using a 20 ppm precursor ion tolerance and requiring each peptides N-C- termini to have trypsin protease specificity, while allowing up to three missed cleavages. TMT tags on peptide N termini/lysine residues (+229.162932 Da) and carbamidomethylation of cysteine residues (+57.02146 Da) were set as static modifications while methionine oxidation (+15.99492 Da) was set as variable modification. A MS2 spectra assignment false discovery rate (FDR) of less than 1% was achieved by applying the target-decoy database search strategy (*107*). Filtering was performed using an in-house linear discrimination analysis algorithm to create one combined filter parameter from the following peptide ion and MS2 spectra metrics: Sequest parameters XCorr and ΔCn, peptide ion mass accuracy and charge state, peptide length and mis-cleavages. Linear discrimination scores were used to assign probabilities to each MS2 spectrum for being assigned correctly and these probabilities were further used to filter the dataset to a 1% protein-level false discovery rate (*104*).

TMT reporter ion and quantitative data analysis. For quantification, a 0.003 m/z window centered on the theoretical m/z value of each ten reporter ions and the closest signal intensity from the theoretical m/z value was recorded. Reporter ion intensities were further de-normalized based on their ion accumulation time for each MS3 spectrum and adjusted based on the overlap of isotopic envelopes of all reporter ions (manufacturer specifications). Total signal to noise values for all peptides were summed for each TMT channel, and all values were adjusted to account for variance in sample handling. For each peptide, a total minimum signal to noise value of 200 was required (*105*, *106*).

#### ChIP-sequencing

The REV-ERBα ChIP-seq experiment was performed as described in (*9*). The data is published GSE79167. In particular, the study was done on BAT samples from three mice acclimated to 29°C harvested at 5 PM (ZT10) with or without a 6 h cold challenge, exactly as previously described in (*9*). The data is published GSE79167. Murine BAT was harvested immediately after euthanasia. It was quickly minced and cross-linked in 1% formaldehyde for 20 min, followed by quenching with 1/20 volume of 2.5 M glycine solution and two washes with ice-cold PBS. Chromatin fragmentation was performed by sonication in ChIP SDS lysis buffer (50 mM HEPES, 1% SDS, 10 mM EDTA at pH 7.5) using probe sonication. Proteins were immunoprecipitated in ChIP dilution buffer (50 mM HEPES, 155 mM NaCl, 1.1% Triton X-100, 0.11% Na-deoxycholate, Complete protease inhibitor tablet at pH 7.5) using the Cell Signaling Technology antibody (#2124). Cross-linking was reversed overnight at 65°C in elution buffer (50 mM Tris-HCL, 10 mM EDTA, 1% SDS at pH 8), and DNA was isolated using phenol/chloroform/isoamyl alcohol. Deep sequencing was carried out by the Functional Genomics Core (J. Schug and K. Kaestner) of the Penn Institute for Diabetes, Obesity, and Metabolism using the Illumina Genome Analyzer IIx and Illumina HiSeq 2000 and sequences were obtained using the Solexa Analysis Pipeline.

### GENE EXPRESSION ANALYSIS

#### RNA extraction and qPCR

Tissues were isolated and immediately snap-frozen in liquid nitrogen. Cells were immediately lysed using TRI reagent (T9424, Sigma-Aldrich). Total RNA was extracted from the tissues and cells using TRI reagent (T9424, Sigma-Aldrich) followed by isolation using RNeasy Mini Kit (74106, Qiagen). Reverse transcription was carried out on 1000 ng RNA using the High Capacity cDNA Reverse Transcription kit (4368814, Applied Biosystems). Gene expression was determined based on real-time quantitative PCR using SYBR green (PP00259, Primerdesign) using LightCycler 480II (Roche) according to manufacturer’s protocol. The data was analyzed with the ΔΔCT method and normalized to the housekeeping gene 36b4. All primers are listed in the **Table S12**.

#### RNA-sequencing and Analysis of *Pparα* AKO BAT

RNA extraction, sequencing and data analysis of BAT from *Pparα* AKO mice was performed exactly as described in the corresponding section of the method details in (*33*). A table of differentially regulated genes is provided in **Table S5**.

#### RNA-sequencing and Analysis of BAT from Ketogenic Diet Fed Mice

The RNA sequencing and data analysis were performed as described in (*69*). In particular, for the generation of RNA-sequencing libraries, polyadenylated mRNA was isolated from 1 μg of total RNA by incubation with oligo-dT beads and prepared according to manufacturer’s protocol (TrueSeq 2, Illumina). Samples were sequenced on the Illumina HiSeq 1500 platform. Sequencing reads were mapped to the mouse or human reference genome (version mm9 or hg19) using STAR (*108*). Tag directories were generated using HOMER (*109*) and exon reads were counted using iRNA-seq (*110*). Normalization and identification of differentially expressed genes was performed using DESeq2 (*111*). A table of differentially regulated genes is provided in **Table S6**.

#### RNA-sequencing and analysis of cultured brown adipocytes

siRNA mediated knock down in murine brown preadipocytes (BMC) and RNA extraction was carried out as stated above. Isolated RNA was treated with DNase I (NEB-M0303L) by mixing 250 ng of RNA, 8 Units of DNase I and DNase buffer in 10 µl reaction volume per sample and incubation the reaction mix for 15 min at 37 °C. Reaction was quenched by thorough vortexing of the samples and incubation at 85 °C for 5 min.

Total RNA integrity (RIN) was assessed using the TapeStation RNA High Sensitivity (HS) kit (Agilent Technologies). mRNA-seq libraries were prepared using the Universal Plus mRNA-seq library preparation kit with NuQuant (TECAN, Männedorf, Switzerland, Cat. #M01485 V8) according to the manufacturer’s instructions. Library concentrations were quantified using Qubit with the NuQuant quantification method and the 1X dsDNA High Sensitivity Assay Kit (Thermo Fisher Scientific, cat. no. Q33231). Library quality was evaluated using the TapeStation DNA D1000 HS kit (Agilent Technologies). Sequencing was performed on an Illumina NovaSeq 6000 platform using SP 100-cycle v1.5 reagents, generating 52-bp paired-end reads. Libraries were sequenced at a molarity of 2 nM with a 1% PhiX spike-in control.

Pre-processing of sequencing data was performed by the Single-Cell Omics platform at the Novo Nordisk Foundation Center for Basic Metabolic Research. In short, sample-demultiplexed FASTQ files were generated from BCL files using the bcl2fastq software (v. 2.20.0.422), and gene-level count matrices were then generated by the nf-core RNA-seq pipeline (v. 3.5, (*112*)). Specifically, this pipeline first trims the reads using the Trim Galore (v. 0.6.7) software, deduplicates the reads based on UMIs via UMI-tools (v. 1.1.2, (*113*)) before finally mapping the reads to the GRCm38 genome using STAR (v. 2.6.1d, (*108*)) and quantifying the gene counts using the salmon software (v. 1.5.2, (*114*)).

Differential gene expression analysis was similarly performed by the Single-Cell Omics platform at the Novo Nordisk Foundation Center for Basic Metabolic Research. Here, all samples were analyzed together using the EdgeR (v. 3.38.0, (*115*)) and limma (v. 3.52.0, (*116*)) R-packages based on the specified contrasts. Specifically, the differential gene expression analyses were performed using the glmQLFit-function with robust=TRUE. Table of differentially regulated genes is provided in **Table S10**.

#### Identification of proteins temporarily co-regulated with SLC25A34

To identify proteins with expression dynamics similar to SLC25A34 during cold adaptation, we performed hierarchical clustering using the Perseus software platform (version 1.5.5.5, (*117*)). The cold adaptation BAT proteomics dataset was analyzed by calculating the Pearson correlation coefficient between each protein and the expression profile of SLC25A34. Proteins were then ranked according to their correlation values, and the top 50 most similarly temporally regulated proteins were selected for downstream pathway enrichment analysis (**Table S4**).

#### Pathway enrichment analysis

All pathway enrichment analyses were performed using GO enrichment analysis, type: PANTHER Overrepresentation Test (Released 20240807). All mouse genes in the database were used as a reference list, the presented pathways are the top 10 enriched pathways sorted based on the Fold Enrichment score with FDR<0.05 (test type: Fisher’s Exact and correction: FDR).

#### Transcription Factor Prediction Based on Differential Gene Expression

To identify potential transcriptional regulators in the dataset, we calculated an average log fold change (logFC) for each gene based on two independent siRNA knockdown datasets (Sigma and Dharmacon) for all genes significantly dysregulated in both datasets (FDR<0,05). Two groups of genes were selected: downregulated genes (logFC ≤ –0.5, n = 542) and upregulated genes (logFC ≥ 0.5, n = 676). These gene groups were used to predict candidate transcription factors using ChEA3 (*118*). The top predicted transcription factors were identified for each group. To contextualize these findings, each transcription factor was annotated based on its expression level in brown adipocytes as high, medium, low, or not detected [ND]), and whether its own mRNA was significantly downregulated following the *Slc25a34* knockdown. The relationship between the transcription factors and their target genes was visualized using clustergrams. Genes forming the clusters indicated on Fig. 5F, S5C are shown in **Table S11**.

### METABOLOMIC ANALYSES

#### Derivatization of extracted metabolites and untargeted metabolomics analyses (**Fig. 4A**, 4B)

siRNA mediated knock down in murine brown preadipocytes (BMC) was carried out as stated above. Cells were washed briefly with ice-cold 1×PBS, and residual PBS was removed by aspiration. Metabolites were extracted by adding 200 μL of ice-cold extraction solvent (methanol:acetonitrile:H_2_0 50:30:20) directly to the cells, followed by scra ping and transfer into pre-labelled, chilled 1.5 mL Eppendorf tubes. Samples were vortexed vigorously for 2 minutes at 4 °C and centrifuged at 1,000 × g for 2 minutes at 4 °C. The resulting supernatant was transferred to new pre-labelled tubes and kept on an ice bath until further processing. For 3-NPH derivatization, samples (20 μL) were derivatized with 50 mM 3-NPH (in 50% acetonitrile) in the presence of 30 mM EDC (in 50% acetonitrile and 6% pyridine) for 45 minutes at 4 °C with gentle shaking. Appropriate blanks, quality control (QC - pool of all samples) and controls were included, including pure standards and samples without extract (solvent only). Derivatized metabolites were analysed by LC-MS as follows. Samples of 3 μl were injected into a Vanquish Horizon UPLC system (Thermo Fisher Scientific) equipped with a Zorbax Eclipse Plus C18 guard column (2.1 × 50 mm, 1.8 μm; Agilent Technologies) and an analytical column (2.1 × 150 mm, 1.8 μm; Agilent Technologies), maintained at 40 °C. Analytes were eluted at a flow rate of 400 μl/min using a gradient of eluent A (0.1% formic acid in water) and eluent B (0.1% formic acid in acetonitrile): 0–1 min, 10% B; 1–1.5 min, 10–40% B; 1.5–5 min, 40–95% B; 5–7.6 min, 95% B; 7.6–8 min, 95–10% B. The system was equilibrated for 3.5 min under initial conditions. The eluent flow was coupled to a TimsTOF Pro (Bruker) for mass spectrometric analysis in negative ion mode. Data were processed using Metaboscape (Bruker, v. 2025) with the In-Silico Derivatization workflow, which includes automated library structure derivatization, CCS prediction, and in-silico fragmentation for automated annotation.

Underivatized samples were extracted as described above. Instead of derivatization, the supernatant was lyophilized and resuspended in 25 μl of 0.1% formic acid in water just prior to analysis. From each sample, 5 μl was transferred to a pooled quality control (QC) sample. From each sample, 2 μl was injected into a Vanquish Horizon UPLC system (Thermo Fisher Scientific) equipped with a Zorbax Eclipse Plus C18 guard column (2.1 × 50 mm, 1.8 μm; Agilent Technologies) and an analytical column (2.1 × 150 mm, 1.8 μm; Agilent Technologies), maintained at 40 °C. Analytes were eluted at a flow rate of 400 μl/min using a gradient of eluent A (0.1% formic acid in water) and eluent B (0.1% formic acid in acetonitrile): 0–1 min, 3% B; 1–3 min, 3–40% B; 3–5 min, 40–95% B; 5–7.6 min, 95% B; 7.6–8 min, 95–3% B. The system was equilibrated for 3.5 min under initial conditions. The eluent flow was coupled to a TimsTOF Pro (Bruker) for mass spectrometric analysis in positive and negative ion modes. Data were processed using Metaboscape (Bruker, v. 2025), with features annotated using the NIST2020 and MetaboBase libraries, integrated as add-ons within Metaboscape. Data provided in **Table S7**.

#### Targeted metabolomic measurement of Acetyl-CoA in cultured brown adipocytes (**Fig. 4C**)

siRNA mediated knock down in murine brown preadipocytes (BMC) was carried out as stated above. Cells were washed briefly with ice-cold 1×PBS, and residual PBS was removed by aspiration. Metabolites were extracted by adding 200 μL of ice-cold extraction solvent (methanol:acetonitrile:H_2_0 50:30:20) directly to the cells, followed by scraping and transfer into pre-labelled, chilled 1.5 mL Eppendorf tubes. Samples were vortexed vigorously for 20 minutes at 4 °C and centrifuged at 1,000 × g for 2 minutes at 4 °C. The supernatant was transferred to a new pre-labbeld tube and lyophilized by vacuum centrifugation. Samples were resuspended in 25 μ of which 3 μl were injected into a Vanquish Horizon UPLC system (Thermo Fisher Scientific) equipped with a Zorbax Eclipse Plus C18 guard column (2.1 × 50 mm, 1.8 μm; Agilent Technologies) and an analytical column (2.1 × 150 mm, 1.8 μm; Agilent Technologies), maintained at 40 °C. Analytes were eluted at a flow rate of 500 μl/min using a gradient of eluent A (50 mM ammonium acetate in water) and eluent B (100% methanol): 0–0.5 min, 10% B; 0.5–7.2 min, 10–99% B; 7.2–8 min, 99% B; 8–8.1 min, 99–10% B; 8.1–12 min, 10% B. The eluent flow was coupled to a Q Exactive HF (Thermo Fisher Scientific) for mass spectrometric analysis in negative ion mode. Data were processed using MZmine 2.53 (*119*) with the Targeted Feature Detection module, targeting CoA esters annotated using in-house standards. Data from the above analyses were processed in Metabolink (*120*) to perform blank subtraction, QC-based filtering, drift correction, data merging, and probabilistic quotient normalization.

#### Untargeted metabolomic analysis of cultured brown adipocytes (**Fig. 4E**, 4G-4K, S4A) Metabolomic Sample Preparation

Metabolites were extracted using a method drafted from (*121*). Briefly, media were aspirated and cells were extracted with 0.7 mL of ice cold 50% aqueous liquid chromatography-mass spectrometry (LCMS) grade acetonitrile (extraction and re-suspension solvent). The plate was then placed on top of wet ice and allowed to rest for 10 minutes.

Cells were scraped in the extraction solvent and transferred to 1.5 mL centrifuge tubes. Samples were centrifuged at top speed for ten minutes in a centrifuge cooled at 0 °C. Blanks were composed by treating an empty well as well as a cell culture well containing media sans cells in the same way as samples. Supernatants were transferred to fresh 1.5 mL centrifuge tubes and dried via speed vacuum. Dried residues were re-suspended in 0.2 mL of 50% aqueous LCMS grade acetonitrile. For the time course study, samples were re-suspended in 0.1 mL of 50% aqueous LCMS grade acetonitrile. All samples were centrifuged for 5 minutes at 20,238 x g without temperature control. 50 µL of supernatant were transferred to fresh autosampler vials. 5 µL of each sample were combined to form a QC pool sample, which was injected 6 times in the beginning of the queue, twice in the middle of the queue and twice at the end of the queue to serve as a performance control. In addition to the above blanks, a solvent blank was composed with the re-suspension solvent. Samples, blanks, and QC pool were placed in a pre-chilled (6 °C) Thermo Scientific Dionex 3000 autosampler and injected as described below. In all analyses, samples were injected in a random order.

#### Liquid Chromatography – Mass Spectrometry

In the non-time course study, samples were injected at 5 µL in negative ion mode. For positive ion mode analysis, samples were diluted 1/10^th^ and injected at 2.5 µL. In the time course study, samples were injected at 5 µL in both modes. Analytes were separated using a gradient at a flow rate of 0.4 mL/min between 0.006% (v/v) LCMS grade ammonium hydroxide in LCMS grade water (mobile phase B) and 0.006% (v/v) LCMS grade ammonium hydroxide (mobile phase B) in LCMS grade acetonitrile over a heated (25 °C) Phenomenex Amide (2 x 100 mm, particle size 3 µm, pore size 100 Å) column fitted with a Phenomenex SecurityGuard guard cartridge containing amide resin. The column was equilibrated at 10% mobile phase B which was then increased to 70% B over ten minutes. The column was washed over five minutes at 100% B, then re-equilibrated at 10% B for ten minutes. Analytes were electrospray ionized and detected in both negative and positive ion mode using a Bruker Impact II Q-TOF mass spectrometer operated in full scan mode (*m/z*: 50 – 1000). The instrument was mass calibrated prior to use on each day using an infusion of 4.5 mM sodium acetate. The calibrant was also infused at the beginning of each analysis and used to mass calibrate each sample. The instrument parameters are as follows: cone voltage, 30 kV; collision energy, 7 eV; nebulizer gas flow, 10 L/min; nebulizer gas temperature, 220°C; nebulizer pressure, 2 bar; and scan rate, 2 Hz. Ions were assigned identities based upon MS/MS fragmentation and matching to Metlin (*122*) and HMDB (*123*) mass spectral libraries.

#### Data Analysis

Raw data was converted to .cdf format using Bruker DataAnalysis software (version 4.3). Peaks were detected, aligned, and integrated using XCMS online (*124*). Ions eluting prior to 1 minute and isotopes were removed from further analysis as well as ions with an intensity below 10,000 counts. MetaboAnalyst (version 4.0) was used for all statistical analyses and heatmap generation (*125*). Outliers were based upon interquartile range. Data were normalized to sum intensity, generalized log-transformed (*126*) and Pareto scaled. All multiple comparisons were false discovery rate (FDR<0.05) corrected. Data are shown in **Table S8** (statistical analysis of the 54 dysregulated metabolites) and **Table S9** (shows relative intensities for detected annotated metabolites that are significantly dysregulated by *Slc25a34* KD at either time point, some metabolites are duplicated because they were detected in both ion modes.

#### Mitochondrial immunoprecipitation and targeted metabolomics analysis of cultured brown adipocytes

##### Cell Culture and Transfection

Immortalized brown preadipocytes were prepared as previously described by . Cells were seeded on 15 cm tissue culture dishes, grown to 100% confluency, and induced to differentiate. On day 3 of differentiation, reverse transfection was performed following the protocol from (*103*). siRNA targeting *Slc25a34* (si*Slc25a34*) or a non-targeting control siRNA (si*Control*) were diluted in Opti-MEM (Gibco, Cat# 51985) to a final siRNA concentration of 300nM. Lipofectamine RNAiMAX (Invitrogen, Cat# 13778-150) was added at a final concentration of 30μL/mL. The siRNA-transfection mix was incubated at room temperature for 30 minutes. In parallel, cells were trypsinized, counted, and resuspended in complete culture medium. A total of 1.5*106 cells per well were seeded on top of the siRNA mixture, resulting in a final siRNA concentration of 50 nM. In addition, adenovirus encoding 3XHA-EGFP-OMP25 (for rapid mitochondrial purification (*47*)) was added dropwise to each well. The media was refreshed two days post-transfection. On day 7 of differentiation, cells were harvested for anti-HA bead pulldown as described in (*128*)

##### Rapid Mitochondria Isolation (HA Pulldown)

Cells were washed twice with PBS and once with 1 mL KPBS. Homogenization was performed using a Dounce homogenizer with 25 strokes. A 50 μL aliquot of the homogenate was collected for western blotting analysis. The remaining homogenate was transferred to a 2 mL microcentrifuge tube and centrifuged at 1000g for 2 minutes. The supernatant was transferred to a fresh 2 mL tube, and 60μL of pre-washed anti-HA magnetic bead slurry (Thermo Scientific, Cat# 88836) was added. Samples were rotated for 3 minutes at 4°C. Beads were magnetically separated, and the flowthrough was collected as the post-mitochondrial (cytosolic) fraction. Beads were then resuspended in 1 mL KPBS and divided into two 500 μL aliquots for downstream metabolomic and western blotting analyses.

##### Metabolite Extraction and Derivatization

Metabolites were extracted by eluting from beads in 380μL of 50% methanol. Samples were centrifuged at 17,000g for 10 minutes, and the supernatants were collected. Supernatants were vortexed for 10 seconds, followed by the addition of 220 μL acetonitrile and another 10-second vortex. Samples were bead-beaten for 6 minutes at 20 Hz using 1.4 mm ceramic beads. Next, 600μL dichloromethane and 300μL water were added, and samples were vortexed for 1 minute before incubating on ice for 10 minutes. Samples were centrifuged for 10 minutes at 1,500g at 1°C, and the aqueous phase was transferred to a new 1.5 mL microcentrifuge tube. To quantify unstable α/β-keto acids (such as oxaloacetate), a reduction was performed prior to sample drying and derivatization as described (*129*). The pH of each aqueous sample was adjusted to 10–12 using 2 M NaOH, verified with pH paper. While on ice, 15 μL of freshly prepared NaBD₄ solution (5 mg/mL in 50 mM NaOH) was added. The reaction was allowed to proceed at room temperature for 1 hour. Samples were returned to ice, and 10μL of cold 1M HCl was slowly added to quench the reaction. Samples were dried using a SpeedVac concentrator.

Dried samples were reconstituted in 30μL of pyridine and then centrifuged at room temperature at 13,000g for 2 minutes. The resulting supernatants were transferred to GC/MS autosampler vials containing 70μL of N-tert-butyldimethylsilyl-N-methyltrifluoroacetamide (MTBSTFA) for derivatization, and then were incubated at 70°C for 1 hour. 1μL of each derivatized sample was injected for GC/MS analysis.

GC/MS analyses were performed using an Agilent 7890A gas chromatograph coupled to a 5975C mass selective detector, equipped with an Agilent 7693 autosampler. Chromatographic separation was achieved using a DB-5ms column (30m length with 10m guard length, 0.25mm inner diameter, 0.25μm film thickness). The GC oven program was initiated with a 1-minute hold at 120 °C, followed by a temperature ramp of 10°C per minute to 300 °C. A final bake-out step was held at 320 °C for 10 minutes. The injector and MS interface temperatures were maintained at 285 °C. Helium was used as the carrier gas at a constant flow rate of 1.5 mL/min. The mass spectrometer operated in electron ionization mode at 70 eV. Data acquisition was performed in selected ion monitoring (SIM) mode and full scan mode (mass range: 50–700 Da). Data analysis was performed using Agilent GCMS MassHunter. Metabolite relative abundance was normalized to protein from each corresponding sample. Data provided in **Table S8.**

### RESPIROMETRY

#### Seahorse XF96 respirometry with intact brown adipocytes

Cells were cultured on Seahorse XF96 Cell Culture Microplates (Agilent Technologies) in the amount of 60000 cells/well. Cell culture medium was changed 1 h before the first measurement to DMEM (Sigma-Aldrich, D5030, supplemented with 143 mM NaCl, pH 7.4) supplemented with 25 mM glucose and 1 mM pyruvate (Gibco, 11360-070) and 4 mM glutamine. Real-time oxygen consumption rate (OCR) was measured under basal conditions and following injections of oligomycin A (1 μM) (Leak) (Cayman Chemicals, 11342), NE (1 μM) (Sigma-Aldrich, A9512) (NE), FCCP (0.5 μM) (Cayman Chemicals, 15218) (MAX), antimycin A/rotenone (1 μM each) (Cayman Chemicals, N/A) (Cayman Chemicals, 13995) (non-mitochondrial oxygen consumption). OCR was assessed using a Seahorse XFe96 Extracellular Flux Analyzer (Agilent Technologies). The presented values represent the measurement values with subtracted non-mitochondrial oxygen consumption) and have not been normalized. The quantification for statistical analysis represents the values of the final measurement for each drug administration (with the exception of NE-stimulated respiration which represents the maximal NE-induced OCR) with subtracted non-mitochondrial oxygen consumption.

#### Seahorse XF96 respirometry with permeabilized brown adipocytes

Cells were cultured on Seahorse XF96 Cell Culture Microplates (Agilent Technologies) in the amount 60000 cells/well. Prior to the experiment, the cells were rinsed with the MiR05 media (see O2k respirometry) once and supplemented with MiR05 media containing the Seahorse XF Plasma Membrane Permeabilizer (Agilent Technologies) in 1 nM concentration, 2 mM ADP (Jena bioscience: NU-1198) and 1 mM MgCl_2_. Immediately after the measurements were started. The compounds were injected in the following concentrations: malate (2 mM) (Sigma-Aldrich: M1000), pyruvate (5 mM) (Sigma-Aldrich: P2256), glutamate (10 mM), succinate (10 mM) (Sigma-Aldrich: S2378), octanoylcarnitine (0.5 mM) (APExBIO Technology: B6371-50), palmitoylcarnitine (0.04 mM) (Sigma-Aldrich: P4509), glycerophosphate (10mM) (Santa Cruz Biotechnology: sc-215789), oligomycin A (1 μM) (Cayman Chemicals, 11342), FCCP (1 μM) (Cayman Chemicals, 15218), antimycin A/rotenone (1 μM) (Cayman Chemicals, N/A) (Cayman Chemicals, 13995). The presented values represent the raw measurement values and have not been normalized. The quantification for statistical analysis represents the maximal OCR values achieved after the respiratory substrate injection without FCCP.

#### *De novo* Lipogenesis Assay in Cultured Mouse Brown Adipocytes

##### Radiolabeling with [¹⁴C]-Glucose

Murine brown adipocytes were cultured and siRNA mediated knockdown was performed as described above in 12 well plates. On day 7 of differentiation, culture media was exchanged to 0.5 mL per well of standard DMEM media supplemented with T3 and Insulin, containing ^14^C-labelled glucose (D-[^14^C(U)] Revvity Health Sciences Inc NEC042V250UC, Lot: 3106342; 250 uCi(9,25 MBq), 265 mCi/mmol (9,805 GBq/mmol), 1,25 ml Ethanol:water 3:97), 0,5 μCi per 0.6 mL of media. A few wells were left untreated as a negative control. Cells were cultivated in the presence of ^14^C-labelled glucose for 6 hours. After 6 hours, cells were rinsed twice with ice-cold PBS and frozen on plates at −70°C until subsequent analysis.

##### Lipid extraction and Scintillation Counting

Lipid extraction was performed as described in (*130*) with modifications. 0.4 mL of H₂O was added to each well and cells were harvested using a rubber policeman into 15 mL conical Falcon tubes. 0.5 mL chloroform and 1.0 mL methanol were added to each sample, followed by vortexing. Next, 0.5 mL chloroform and 0.5 mL of 0.2 N HCl were added, and samples were vortexed again. Phase separation was achieved by centrifugation at 2500 rpm for 5 min at room temperature. The lower (organic) phase was transferred to a clean 15 mL tube and washed once with a pre-prepared “blank upper phase” (generated by mixing 1 mL chloroform, 1.0 mL methanol, 0.5 mL of 0.2 N HCl, and 0.4 mL water per sample). After a second centrifugation, the cleaned lower phase was transferred to a 2 mL glass vial and organic solvents were evaporated to dryness under nitrogen flow. Dried lipids were resuspended in 75 µL chloroform:methanol (2:1, v/v), and 60 µL was transferred to scintillation vials containing 5 mL scintillation fluid (Ultima Gold, Revvity Health Sciences Inc 6013329). Scintillation was counted using Hidex 600 SLe Automatic TDCR Liquid Scintillation Counter.

##### Citrate Synthase Activity

The mitochondria abundance in the cultured brown adipocytes (BMC) was estimated by measuring the activity of citrate synthase (CS) in the cellular extracts. The assay is based on the protocol from (*131*) and was performed as described in (*129*) with modifications. Briefly, the cells were grown in 6 well plates (1000000 cells per well) and harvested by adding 400 μl of the lysis buffer (50 mmol/l Hepes, 137 mmol/l NaCl, 10 mmol/l Na4P2O7, 20 mmol/l NaF, 5 mmol/l EDTA, 1 mmol/l MgCl2, 1 mmol/l CaCl2, 2 mmol/l Na3VO4, 5 mmol/l nicotinamide, 10 µmol/l trichostatin A, Halt protease inhibitor cocktail (Thermo Scientific, Waltham, MA, USA), 1% (v/v) NP-40, 10% (v/v) glycerol, demineralized water). The extracts were mixed with reaction mix containing 0.1 m Tris-HCl, pH 8.1, 0.4 mm acetyl-CoA sodium salt (Sigma-Aldrich) and 1 µm 5,5′-dithiobis(2-nitrobenzoic acid) (Sigma-Aldrich), and absorbance at 412 nm was measured every 30 s for 5 min (basal slope). Oxaloacetic acid at 10 mm was used as the reaction substrate, and the change in absorbance after its addition to the mix was measured every 30 s for another 5 min (reaction slope). CS activity was then calculated from the delta of the reaction and basal slopes and presented in µmol/min/g protein. Protein content was measured in a separate set of samples produced in the same experiment using Bradford assay (Quick Start™ Bradford 1x Dye Reagent #5000205m Bio-Rad) according to manufacturer protocol.

### MOLECULAR DOCKING

The hydrated and flexible docking simulations were performed in Python using Autodock Vina (*133*, *134*). AlphaFold PDB file containing the receptor coordinates for Slc25a34 (AF-Q6PIV7-F1-v4) were first modified with the reduce script (*135*) ($ reduce -BUILD receptor.pdb), adding all missing hydrogen atoms. PDBQT file and grid parameters of the receptor were then generated using the python package Meeko ($ mk_prepare_receptor.py) with the following grid coordinates (x y z ; box size), 0 0 0; 30 30 30 (*136*). E28, K31, K128, R176, D228 and R277 were specified to be flexible residues. Affinity maps and water maps were finally computed with autogrid4 and the python Vina package mapwater.py, respectively. Ligand coordinates were obtained through PubChem (sdf files, 3D structures) and hydrogen atoms were first added with the python package Molscrub ($ scrub.py -I ligand.sdf -0 ligandH.sdf). The meeko script ($ mk_prepare_ligand.py -i ligandH.sdf -o ligand.pdbqt -w) was then used to add explicit water molecules to the ligands and generate the PDBQT files (*136*). The Docking affinities were calculated using Autodock Vina with Autodock 4 forcefield and the exhaustiveness set at 32, following Vina hydrated docking protocol (https://autodock-vina.readthedocs.io/). The computed docking were visualized in the software PyMOL (Schrodringer), showing residues implicated in the matrix salt bridges of the transporter, as well as interacting with the ligands.

### QUANTIFICATION AND STATISTICAL ANALYSIS

All statistical tests were performed using GraphPad Prism software. Data are presented as means+SEM. All statistical tests are stated in the figure legends. For the Two-way ANOVA tests corrections for multiple comparisons were made by controlling the False Discovery Rate (FDR) using the Two-stage step-up method of Benjamini, Krieger, and Yekutieli unless otherwise indicated in the figure legends.

## SUPPLEMENTARY FIGURE LEGENDS

**Figure S1:**
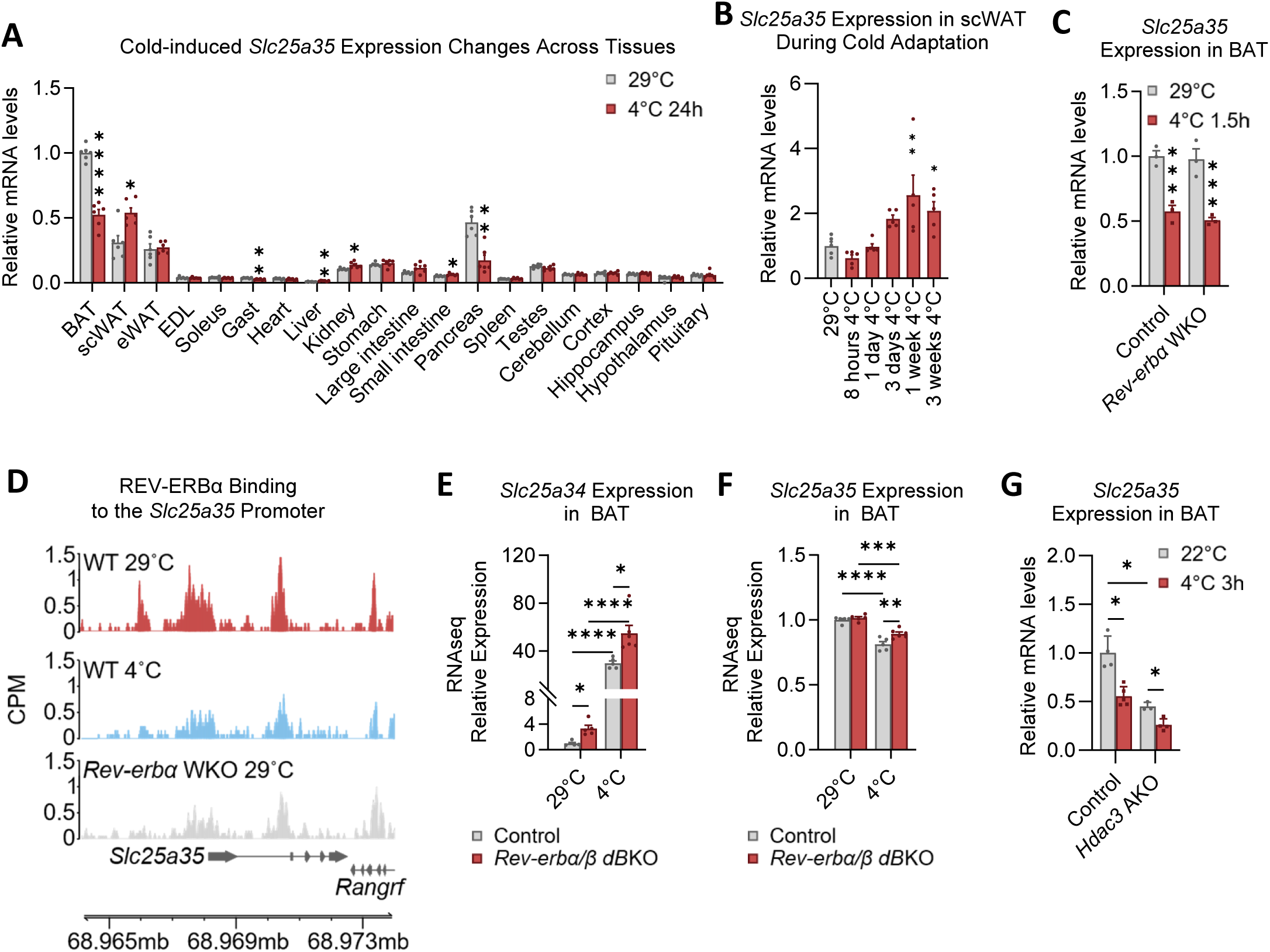
**A**, *Slc25a35* expression across tissues, fold change relative to the expression level in the tissue at TN. **B**, *Slc25a35* mRNA levels in scWAT during cold adaptation. **C**, *Slc25a35* mRNA levels in BAT of control and *Rev-erbα* KO mice at TN and cold. **D**, *Slc25a35* mRNA levels in BAT from control and *Hdac3* AKO mice at TN and after 6 h of cold exposure (ZT10). For all panels, error bars represent ±SEM, p < 0.05 = *, p < 0.01 = **, p < 0.001 = ***, p < 0.0001 = ****, unpaired two-tailed multiple t-tests (A), One-way ANOVA (B) and Two-way ANOVA (C, D) with the Benjamini, Krieger, and Yekutieli two-stage linear step-up procedure for multiple comparisons correction.

**Figure S2:**
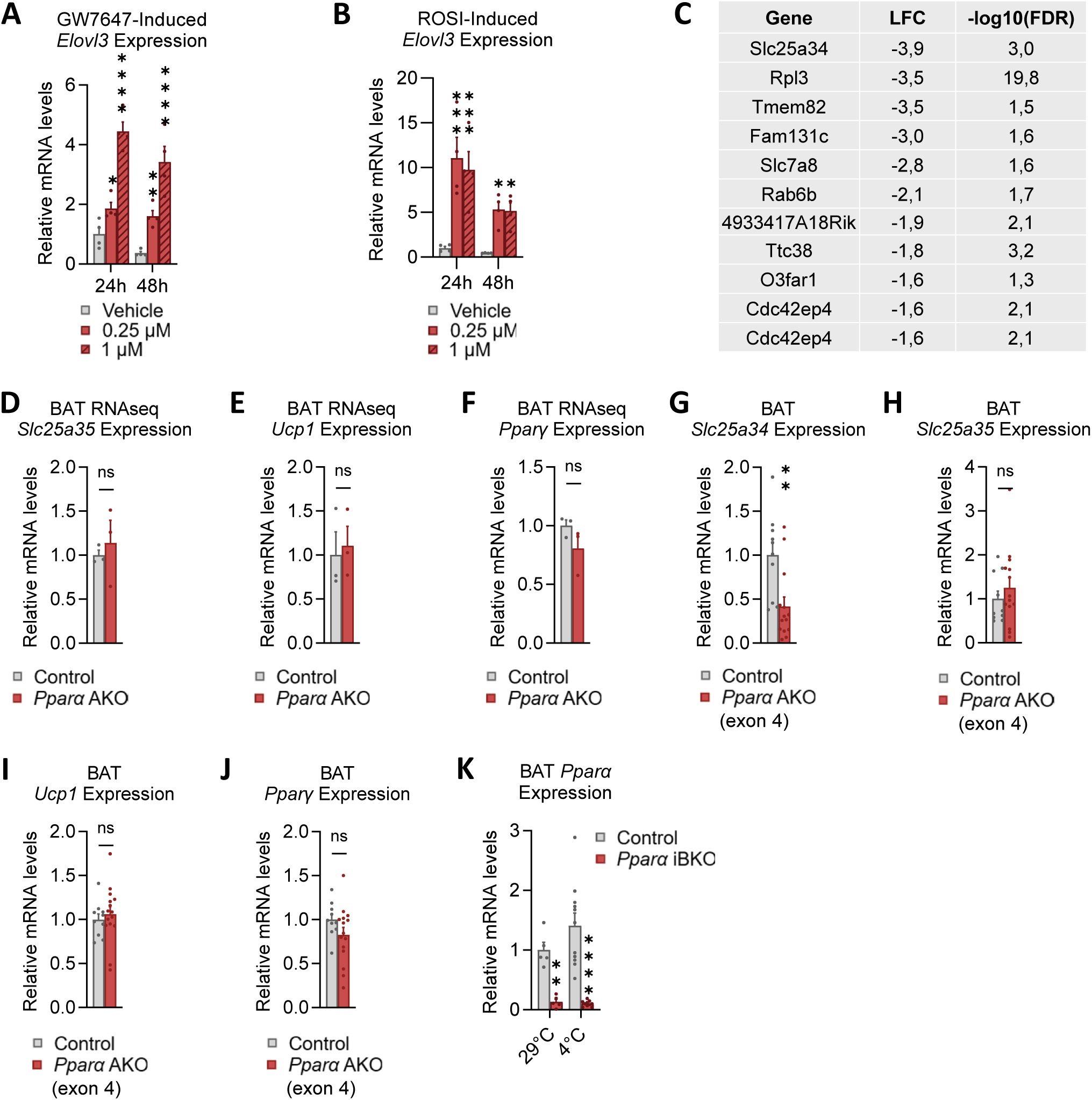
Regulation of *Elovl3* mRNA by, **A**, PPARα agonist GW7647 or, **B**, PPARγ agonist rosiglitazone in cultured brown adipocytes. **C**, Genes significantly decreased (LFC<1, FDR<0.05) in *Pparα* AKO (targeted exon 5) BAT compared to control littermates. **D-F**, Gene expression in BAT of control and *Pparα* AKO (targeted exon 5) mice quantified in RNA-seq analysis. **G-J**, Gene expression in BAT of control and *Pparα* AKO (targeted exon 4) mice. **K**, *Pparα* gene expression in BAT from control and *Pparα* iBKO mice. For all panels, data are represented as mean ±SEM, p < 0.05 = *, p < 0.01 = **, p < 0.001 = ***, p < 0.0001 = ****, Two-way ANOVA (A, B) with the Benjamini, Krieger, and Yekutieli two-stage linear step-up procedure for multiple comparisons correction, for (D-F) p-value was computed using Bioconductor software’s package and corrected for multiple testing using the Benjamini & Hochberg mode of the R function *p.adjust* to compute a false discovery rate (FDR), (G-J) unpaired two-tailed multiple t-tests, (K) Two-way ANOVA with the Benjamini, Krieger, and Yekutieli two-stage linear step-up procedure for multiple comparisons correction.

**Figure S3:**
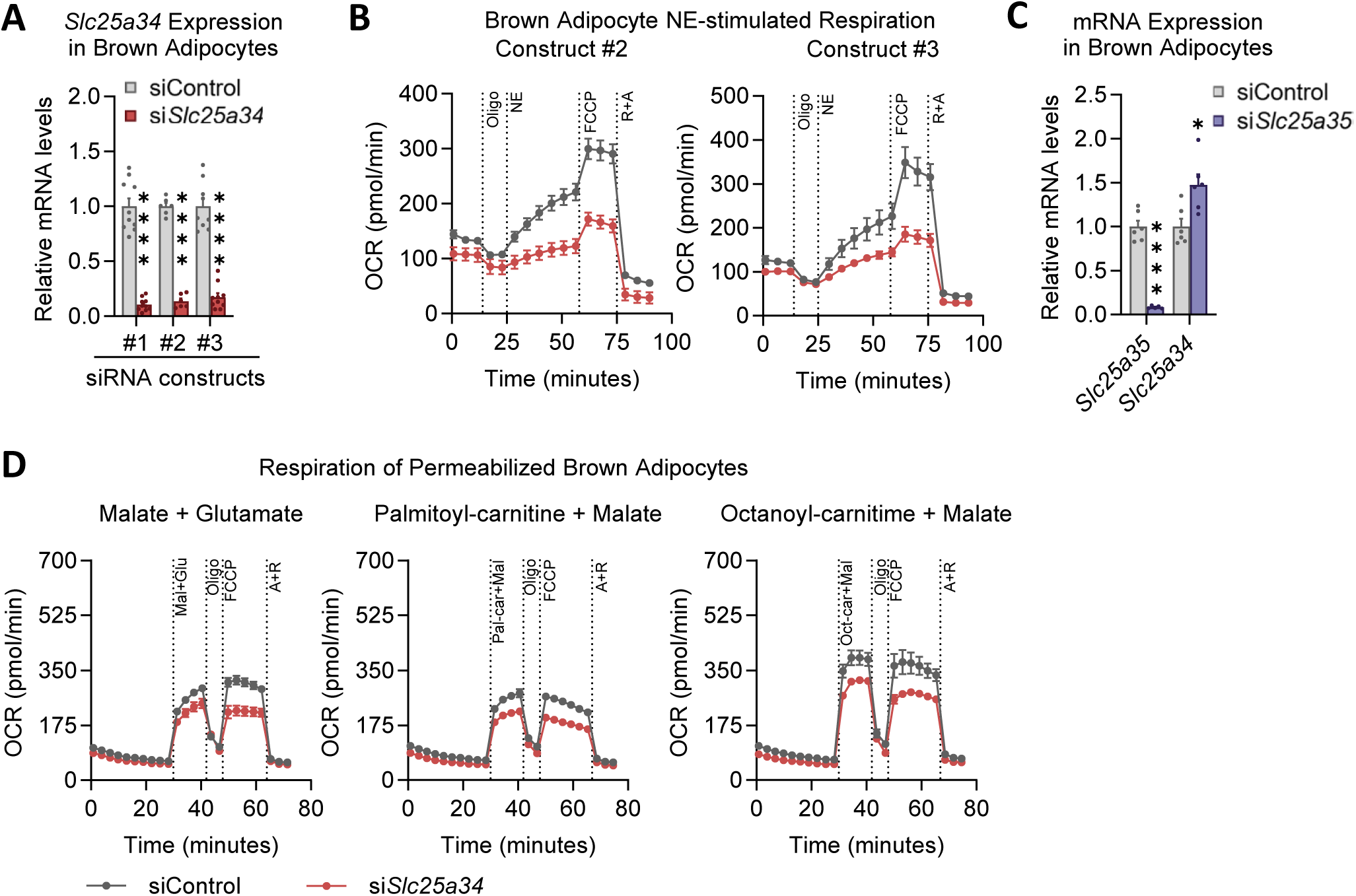
**A**, Validation of siRNA-mediated *Slc25a34* knockdown from three siRNAs from two vendors (Construct #1, Construct #2, Construct #3). **B**, NE-induced oxygen consumption in brown adipocytes with control or siRNA-mediated *Slc25a34* knockdown (Construct #2, Construct #3). **C**, mRNA levels in brown adipocytes with control or siRNA-mediated *Slc25a35* knockdown. **D**, respiration of permeabilized brown adipocytes following 4 days of siRNA-mediated *Slc25a34* knockdown (construct #1) and addition of specific respiratory substrates (from left to right): malate + glutamate, palmitoyl-carnitine + malate, and octanoyl-carnitine + malate. For all panels, data are represented as mean ±SEM, p < 0.05 = *, p < 0.0001 = ****, unpaired two-tailed multiple t-tests) with the Benjamini, Krieger, and Yekutieli two-stage linear step-up procedure for multiple comparisons correction (A, D).

**Figure S4:**
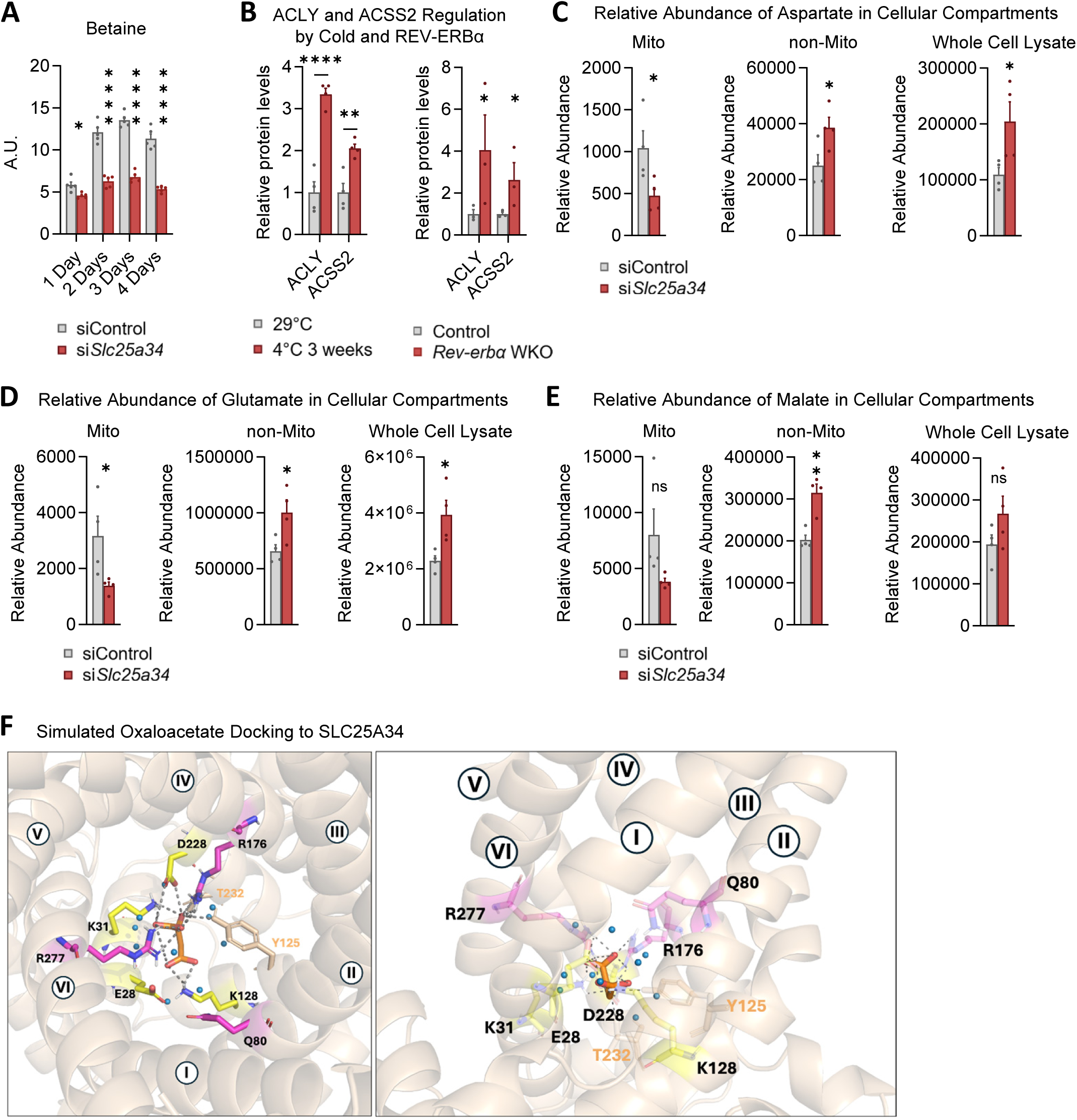
**A**, Changes in betaine levels in an *Slc25a34* siRNA-mediated knockdown time-course in brown adipocytes. **B**, Regulation of ACSS2 and ACLY protein levels by cold in wild type mice (left) and by REV-ERBα in *Rev-erbα* WKO mice (right). **C-E**, abundance of aspartate, glutamate, and malate across cellular compartments. *Slc25a34* knockdown on all panels was performed with siRNA construct #1. **F**, simulated docking of oxaloacetate (OAA) to human SLC25a34 from Alphafold as visualized in Pymol. Residues in the common binding sites are colored pink, residues that form the matrix salt bridge network are colored yellow. Other residues found to form ionic polar interactions with OAA are shown in wheat. Dashed grey lines represent polar interactions. The left and right figure panels provide the cytosolic and lateral view of OAA docking to SLC25A34, respectively. For all panels, data are represented as mean ±SEM, p < 0.05 = *, p < 0.01 = **, p < 0.0001 = ****, Two-way ANOVA with the Benjamini, Krieger, and Yekutieli two-stage linear step-up procedure for multiple comparisons correction (A, B-left), Wilcoxon rank-sum test (B-right), unpaired two-tailed Student’s t-test (C-E).

**Figure S5:**
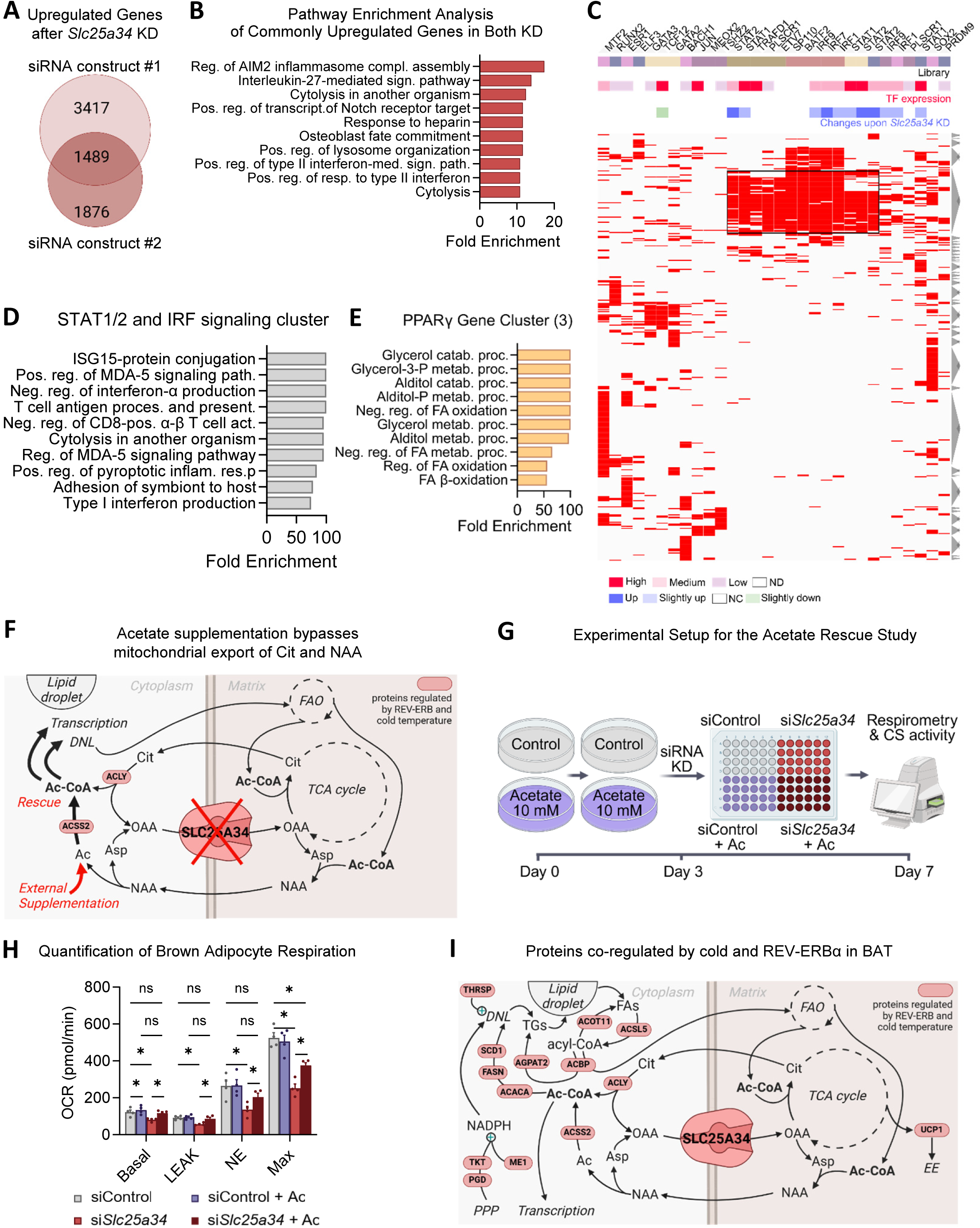
**A**, Comparison of genes significantly upregulated following *Slc25a34* knockdown using two different siRNAs. **B**, Pathway enrichment for the conserved gene signature upregulated by both siRNAs. **C**, Promoter motif analysis of the genes upregulated by loss of *Slc25a34* for candidate transcription factors and, **D**, pathway enrichment of the factor-specific gene cluster. **E**, Pathway enrichment of the PPARγ-specific gene cluster from the genes downregulated by loss of *Slc25a34*. **F**, Model of how acetate supplementation replenishes cytosolic acetyl-CoA production bypassing the mitochondrial export of citrate (Cit) and N-acetylaspartate (NAA). **G**, Schematic showing experimental design of the acetate rescue study. **H**, Effect of acetate supplementation on respiration and mitochondrial content of brown adipocytes with siRNA-mediated Slc25a34 knockdown; quantification of maximal OCR values reached during each run in 4 independent experiments; Slc25a34 knockdown was performed with siRNA Construct #1, measurements done after 4 days of knockdown induction. **I**, Schematic showing BAT proteins co-regulated by cold and the circadian clock and their involvement in numerous lipid metabolism pathways. For all panels, data are represented as mean ±SEM, p < 0.05 = *, p < 0.01 = **, RM two-way ANOVA with the Geisser-Greenhouse correction, matched values are spread across a row with Tukey’s multiple comparisons test, with individual variances computed for each comparison (H).

